# Basal forebrain volume reliably predicts the cortical spread of Alzheimer’s degeneration

**DOI:** 10.1101/676544

**Authors:** Sara Fernández-Cabello, Martin Kronbichler, Koene R. A. Van Dijk, James A. Goodman, R. Nathan Spreng, Taylor W. Schmitz, for the Alzheimer’s Disease Neuroimaging Initiative

**Author notes:** Data used in preparation of this article were obtained from the Alzheimer’s Disease Neuroimaging Initiative (ADNI) database (adni.loni.usc.edu). As such, the investigators within the ADNI contributed to the design and implementation of ADNI and/or provided data but did not participate in analysis or writing of this report. A complete listing of ADNI investigators can be found at: http://adni.loni.usc.edu/wp-content/uploads/how_to_apply/ADNI_Acknowledgement_List.pdf.

## Abstract

Alzheimer’s disease neuropathology is thought to spread across anatomically and functionally connected brain regions. However, the precise sequence of spread remains ambiguous. The prevailing model posits that Alzheimer’s neurodegeneration starts in the entorhinal cortices, before spreading to temporoparietal cortex. Challenging this model, we previously provided evidence that degeneration within the nucleus basalis of Meynert (NbM), a subregion of the basal forebrain heavily populated by cortically projecting cholinergic neurons, precedes and predicts entorhinal degeneration (Schmitz and Spreng, 2016). There have been few systematic attempts at directly comparing staging models using in vivo longitudinal biomarker data, and determining if these comparisons generalize across independent samples. Here we addressed the sequence of pathological staging in Alzheimer’s disease using two independent samples of the Alzheimer’s Disease Neuroimaging Initiative (*N1* = 284; *N2* = 553) with harmonized CSF assays of amyloid (Aβ) and hyperphosphorylated tau (pTau), and longitudinal structural MRI data over two years. We derived measures of gray matter degeneration in a priori NbM and the entorhinal regions of interest. To examine the spreading of degeneration, we used a predictive modelling strategy which tests whether baseline gray matter volume in a seed region accounts for longitudinal change in a target region. We demonstrated that predictive pathological spread favored the NbM→entorhinal over the entorhinal→NbM model. This evidence generalized across the independent samples (*N1*: *r*=0.20, *p*=0.03; *N2*: *r*=0.37, *p*<0.001). We also showed that CSF concentrations of pTau/Aβ moderated the observed predictive relationship, consistent with evidence in rodent models of an underlying trans-synaptic mechanism of pathophysiological spread (*t*_826_=2.55, *p*=0.01). The moderating effect of CSF was robust to additional factors, including clinical diagnosis (*t*_826_=1.65, *p*=0.49). We then applied our predictive modelling strategy to an exploratory whole-brain voxel-wise analysis to examine the spatial specificity of the NbM→entorhinal model. We found that smaller baseline NbM volumes predicted greater degeneration in localized regions of the entorhinal and perirhinal cortices. By contrast, smaller baseline entorhinal volumes predicted degeneration in the medial temporal cortex, recapitulating the prevailing staging model. Our findings suggest that degeneration of the basal forebrain cholinergic projection system is a robust and reliable upstream event of entorhinal and neocortical degeneration, calling into question the prevailing view of Alzheimer’s disease pathogenesis.

## Introduction

Alzheimer’s disease neuropathology progresses in stages, with certain brain regions affected by neuronal deposition of insoluble amyloid (Aβ) and hyper phosphorylated tau (pTau) before others. The asymmetrical progression of Alzheimer’s disease neuropathology in different brain regions may reflect selective neuronal vulnerabilities local to each region (Mattson and Magnus, 2006; Saxena and Caroni, 2011; Wu *et al.*, 2014; Schmitz and Nathan Spreng, 2016). However, evidence in both mouse models of Alzheimer’s disease (De Calignon *et al.*, 2012; Liu *et al.*, 2012) and in human Alzheimer’s disease patients (Schmitz *et al.*, 2018; Sepulcre *et al.*, 2018) indicates that neuropathology also spreads across anatomically and functionally connected brain regions. It is therefore possible that pathologies arising from selective neuronal vulnerability local to a given brain region might precede and predispose the subsequent spread of pathology across networks (Warren *et al.*, 2013). Despite these observations, the precise sequence of neuropathological spread and its relationship with neurodegeneration during the earliest stages of disease remains ambiguous, and can only be resolved with in vivo longitudinal data integrating measures of proteinopathy and microstructural change. Moreover, there have been few systematic attempts at generalizing staging models across independent samples, which limits their clinical utility. Determining when and why certain brain regions are affected in the course of Alzheimer’s disease is essential for understanding the mechanistic basis of disease progression, accurately staging the disease, and identifying individuals for early therapeutic interventions.

The prevailing staging model posits that abnormal Aβ and pTau accumulation starts in the entorhinal cortices (Braak and Braak, 1991; Corder *et al.*, 2000; Braak *et al.*, 2006), before spreading to interconnected areas of medial temporal and posterior parietal cortex. The selective vulnerability of neurons in the entorhinal cortex to these proteinopathies is attributed to their high metabolic activity, the complexity of their axonal projections and the lifelong maintenance of their axonal plasticity (Mattson and Magnus, 2006; Liu *et al.*, 2012; Khan *et al.*, 2014; Wu *et al.*, 2014; Roussarie *et al.*, 2018). One individual spiny stellate neuron in layer II of the entorhinal cortex innervates the entire transverse axis of the dentate gyrus, CA2/CA3 and the subiculum (Tamamaki and Nojyo, 1993). Under a recent framework, amyloid pathology in entorhinal neurons potentiates hyper-phosphorylation of tau proteins; Khan *et al.*, 2014; Wu *et al.*, 2016), which then spread via trans-synaptic mechanisms to distal neocortical areas which receive inputs from EC; i.e. the EC→Neocortical model. See Figure 1A.

**Figure 1.**
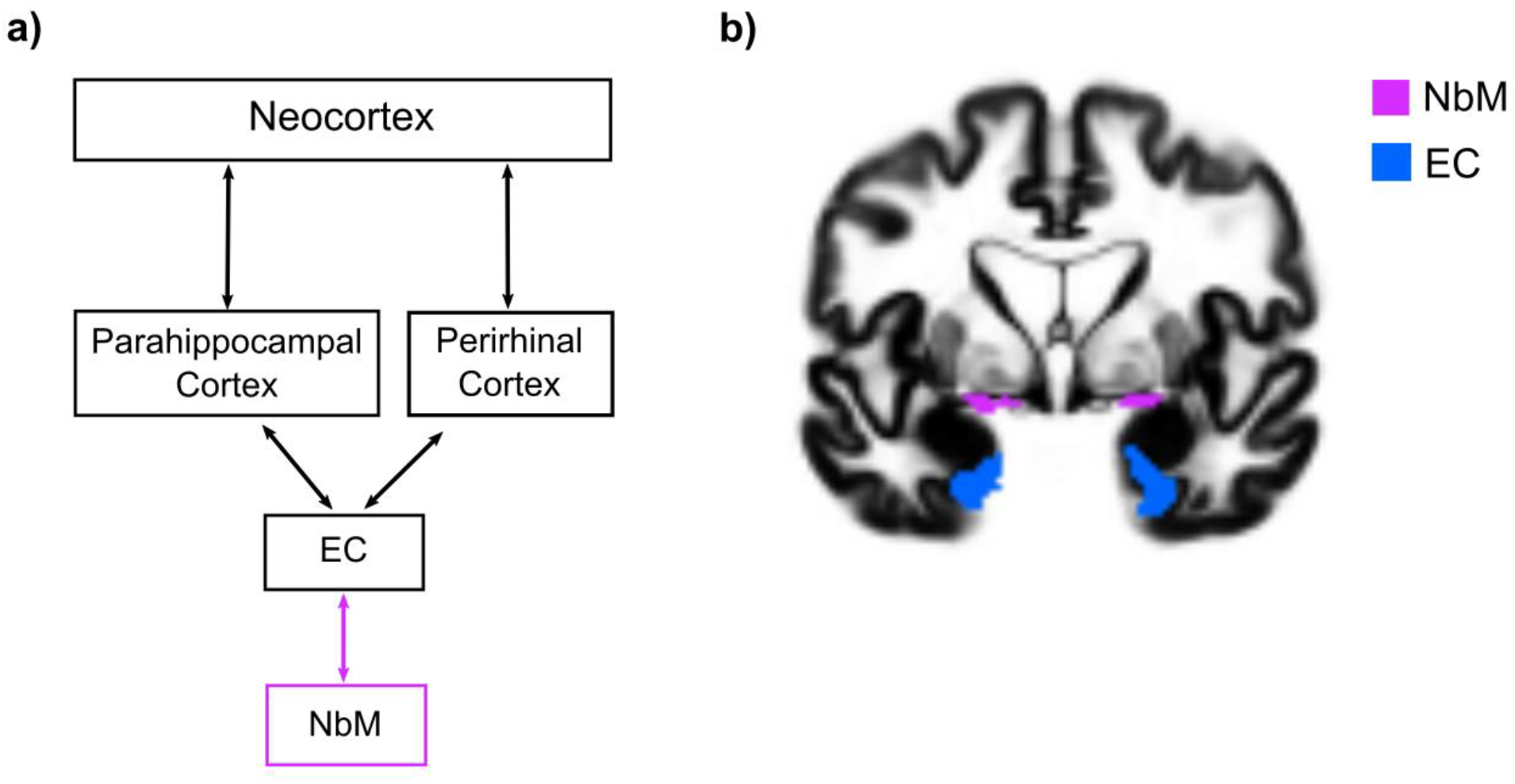
Proposed BF→entorhinal→neocortical model and regions of interest. (a) Predictive pathological staging model of tau pathology originating in EC and spreading via trans-synaptic mechanism to cortical targets of its projection system (black squares) updated to incorporate the BF degeneration as an ‘upstream’ event in the pathological cascade (purple square). (b) Regions of interest of the NbM (purple) and the EC (blue) are superimposed on a coronal slice of the population DARTEL template. Panel (a) is adapted with permission from (Liu *et al.*, 2012).

Challenging the EC→Neocortical model, we have provided initial evidence that degeneration within the nucleus basalis of Meynert (NbM), a subregion of the basal forebrain, precedes and predicts degeneration in the EC (Schmitz and Spreng, 2016). The NbM (also referred to as Ch4 in Mesulam’s nomenclature (Mesulam, 1983; Mesulam and Geula, 1988) is heavily populated (90% of cell bodies) by cortically projecting cholinergic neurons (Mesulam *et al.*, 2004) which are also known for their extremely long and complex branching; the full arborization length of a single neuron was estimated in humans at >100 meters (Wu *et al.*, 2014). Like the layer II entorhinal spiny projection neurons, the cortical cholinergic projection neurons of the basal forebrain accumulate Aβ and pTau early in the course of Alzheimer’s disease, and as early as the third decade of life (Mesulam *et al.*, 2004; Baker-Nigh *et al.*, 2015). Lending further support to early vulnerability of the basal forebrain to Alzheimer’s disease, we previously found that longitudinal NbM degeneration was selectively increased in cognitively normal older adults with abnormal CSF biomarkers of Aβ, over and above age-related degeneration in cortical areas including entorhinal cortex (Schmitz and Spreng, 2016). These initial findings raise the possibility that Alzheimer’s disease pathology spreads from the basal forebrain to entorhinal cortex (NbM→EC), possibly via the same trans-synaptic mechanisms of tau spread proposed previously in the rodent research (Liu *et al.*, 2012). If true, the NbM→EC model would add a crucial ‘upstream’ link to the EC→Neocortical predictive pathological staging of Alzheimer’s disease. Under this model, the ascending cholinergic projections from NbM first ‘seed’ the EC with tau pathology, thereby eventuating the pattern of temporoparietal neurodegeneration typically attributed to the earliest stages of Alzheimer’s disease. Support for the NbM→EC model would have major clinical implications, motivating a shift toward preventative treatment strategies aimed at relieving age-related pressures on the cortical cholinergic projection system.

Here we directly compared the NbM→EC and EC→NbM models of predictive pathological staging in a large sample of older adults ranging from cognitively normal to Alzheimer’s dementia, and then examined if the results generalized to a second independent sample of adults spanning the same clinical diagnostic continuum. In each sample, we first integrated CSF biomarkers of Aβ and pTau neuropathology into a ratio pTau/AB and delineate two groups of older adults in each of the two samples using an independently defined ratio cutpoint which was recently cross-validated at ~90% sensitivity and specificity (Hansson 2018, Schindler 2018). Individuals below the cutpoint exhibit a ‘neurotypical’ biomarker-based phenotype of brain aging; individuals above the cutpoint exhibit an ‘Alzheimer’s pathological’ phenotype. Because CSF pTau/AB is sensitive to the earliest abnormalities of AD (Jack et al 2013), we identified many cognitively normal adults falling above the cutpoint in both samples. We then tracked these groups with structural MRI (sMRI) indices of baseline gray matter volume and longitudinal neurodegeneration over a two-year period. Both the CSF and sMRI data were acquired from multiple phases of the Alzheimer’s disease Neuroimaging Initiative (ADNI).

To examine the ‘spread’ of degeneration between NbM and EC, we used a predictive modelling strategy which tests whether baseline gray matter volume in a seed region accounts for variation in subsequent longitudinal degeneration (change over future timepoints) in a target region. We determined whether evidence of predictive pathological spread favored either the NbM→EC or EC→NbM model, and whether this evidence generalized across the independent samples. We then determined whether CSF concentrations of pTau/Aβ moderated the observed predictive relationship, consistent with an underlying trans-synaptic mechanism of pathophysiological spread. Additional moderating factors, including clinical diagnosis, were evaluated in this framework. We then examined the selectivity of predictive pathological spread between NbM and EC using a whole-brain voxel-wise predictive modelling strategy. Finally, we used the same whole-brain strategy to test whether the EC→Neocortical model recapitulates the spread of neuropathology from EC to temporoparietal cortices observed in prior work (Liu *et al.*, 2012; Khan *et al.*, 2014). If degeneration in the basal forebrain cholinergic projection system is a robust and reliable upstream event of entorhinal degeneration, then it should predict localized entorhinal degeneration, whereas entorhinal cortex should predict downstream events in temporal and parietal cortices.

## Materials and methods

### ADNI Data

Data used in the preparation of this article were obtained from the Alzheimer’s Disease Neuroimaging Initiative (ADNI) database (adni.loni.usc.edu). The ADNI was launched in 2003 as a public-private partnership, led by Principal Investigator Michael W. Weiner, MD. The primary goal of ADNI has been to test whether serial magnetic resonance imaging (MRI), positron emission tomography (PET), other biological markers, and clinical and neuropsychological assessment can be combined to measure the progression of mild cognitive impairment (MCI) and early Alzheimer’s disease (AD).

We used data from the ADNI-1 as first cohort and ADNI-GO and ADNI-2 combined (ADNI-GO/2) as second cohort. ADNI-1 was carried out in 1.5T MR scanners and ADNI-GO/2 in 3T MR scanners. All high-resolution T1 sMRI scans were downloaded from ADNI LONI (http://adni.loni.usc.edu/). A total of 502 ADNI-1 sMRI scans were downloaded from a standardized 2-years interval image collection. This data collection was created to minimize variability between studies and ensure a minimum quality control of the images (Wyman *et al.*, 2014). To obtain the ADNI-GO/2 sMRI scans, we first selected those subjects that had CSF biomarker data (see below) and searched by their research IDs (RIDs). We downloaded longitudinal sMRI images from 714 ADNI-GO/2 participants. To select the two longitudinal time points for each individual, we applied a bounded interval of a mean=1.5 years +/-12 months to maximize the inclusion of participants. 104 participants didn’t have longitudinal sMRI data within the bounded interval. For ADNI sites with GE and Siemens scanners, we used images corrected for distortions (GradWarp) and B1 non-uniformity. For ADNI sites with Philips scanners, these corrections are applied at acquisition (see http://adni.loni.usc.edu/methods/mri-tool/mri-pre-processing/). Finally, to be able to compare ADNI-1 and ADNI-GO/2 data as two independent samples, we discarded participants from ADNI-GO/2 that also took part in the ADNI-1 studies (n=49), due to the smaller sample size of the ADNI1 cohort.

### CSF biomarkers

Accumulation of brain Aβ in plaques and pTau in neurofibrillary tangles are the main neuropathological signs of AD. These biomarkers can be measured in the CSF and are able to differentiate Alzheimer’s disease patients in good concordance with positron emission tomography classifications (Shaw *et al.*, 2009; Hansson *et al.*, 2018; Schindler *et al.*, 2018). CSF samples of Aβ and pTau from the baseline visit were produced with a fully-automated Elecsys protocol (see Supplementary section *CSF procedures*). We calculated a ratio of pTau/Aβ and used a standardized cut-off of 0.028, which was recently cross-validated between the ADNI and Swedish BioFinder studies (Hansson *et al.*, 2018) to classify participants into abnormal (aCSF; pTau/Aβ >= 0.028) and a normal CSF groups (nCSF < 0.028). Using ratios of pTau over Aβ supersedes using single analytes to distinguish amyloid status using positron emission tomography (Schindler *et al.*, 2018). 216 participants from ADNI-1 didn’t have CSF measures and were therefore not included. It is important to note that our grouping strategy is purely based on CSF biomarkers, independent of any cognitive or clinical diagnosis, though we follow-up with analyses of clinical diagnosis (see Results).

### APOE genotyping

The ε4 allele of the APOE gene is the strongest genetic risk factor for non-familial Alzheimer’s disease (Corder *et al.*, 1993; Liu and Bu, 2013) and is thought to interact with the selective vulnerability of cholinergic BF neurons by disrupting the capacity of these cells to support and maintain their enormous axonal membranes (Poirier *et al.*, 1993, 1995; Poirier, 1994). In order to study the effect of the CSF Alzheimer’s disease biomarkers independently of other factors, we included the APOE genotype as a covariate of no interest in our analyses. The APOE information for all individuals with both sMRI and CSF data was obtained from the APOERES.csv spreadsheet and the ε4+ status was defined as having at least one ε4 allele. A blood sample at the screening or baseline visit was collected for APOE genotyping and genotype analysis was performed following standard procedures (Saykin *et al.*, 2010).

### Neuropsychological assessment

Cognitive performance was assessed with a neuropsychological battery that covered a wide range of cognitive functions, including short and long-term memory, language, executive function and attention. We used baseline data from two composite scores, one for memory and one for executive function, derived from the ADNI neuropsychological battery that were recently validated using confirmatory factor analysis (Crane *et al.*, 2012; Gibbons, 2012). Specifically, the memory score integrated the Mini-mental state examination (MMSE), Rey Auditory Verbal Learning Test, Alzheimer’s Disease Assessment Scale, and Logical Memory tests. The executive function score integrated the Category Fluency, Digit Span Backwards, Digit Symbol Substitution, Trails A and B and Clock Drawing tests.

### ADNI Clinical Diagnosis

At the baseline visit, an initial diagnosis recommended by the ADNI Clinical Core was used to classify participants into Cognitively Normal (CDR=0, MMSE 24–30), Mild Cognitively Impaired (CDR=0.5, MMSE 24–30) and Alzheimer’s disease (CDR 0.5-1, MMSE 20-26) (Aisen *et al.*, 2010, 2015; Petersen *et al.*, 2010).

### CSF groups by neuropsychological status

To confirm that our CSF grouping strategy differentiated participants based on measures of neuropsychological status, we collapsed the baseline neuropsychological data from both ADNI cohorts to compare the aCSF and nCSF groups on the Mini Mental State Examination, a common test of general cognitive function, as well as the two composite scores of memory and executive function (see preceding section). For each test, we computed one way fixed effects ANOVAs with CSF group (aCSF, nCSF) as a factor, covarying for Age, Sex, Education, Cohort and APOE genotype. These models revealed significant main effects of CSF pathology on cognition in all tests. Consistent with the sensitivity of CSF pTau/Aβ to pathological aging, the aCSF group exhibited significantly more cognitive impairment than the nCSF group on Mini Mental State Examination, memory and executive function. See Table 1.

**Table 1.** Neuropsychological Function by CSF Group (collapsed across ADNI cohorts)

|  | CSF group |  | F | p |
| --- | --- | --- | --- | --- |
|  | aCSF | nCSF |  |  |
| Mini Mental State Examination | 26.28 (2.83) | 28.65 (1.57) | $F_{1,830}=112.74$ | $p<0.001$ |
| Memory | -0.19 (0.81) | 0.86 (0.68) | $F_{1,829}=243.77$ | $p<0.001$ |
| Executive Function | -0.20 (1.01) | 0.72 (0.88) | $F_{1,829}=123.96$ | $p<0.001$ |

Means and standard deviations of the neuropsychological tests. The aCSF group exhibited worse general cognition (MMSE), memory and executive function compared to the nCSF group. MMSE = Mini Mental State Examination; aCSF = abnormal CSF group; nCSF = normal CSF group.

### Longitudinal structural MRI

A serial longitudinal pipeline scheme in SPM12 r7219 (https://www.fil.ion.ucl.ac.uk/spm/) implemented in Matlab 2013a was used to calculate the GM volume changes from the sMRI data (Ashburner and Ridgway, 2013). This procedure unifies rigid-body registration, intensity inhomogeneity correction and non-linear diffeomorphic transformation in a single iterative model. Initially, all of the individual time-points were co-registered to the ICBM 152 volume to ensure good starting parameters for the high-dimensional DARTEL warping algorithm. We then employed within-subject symmetric diffeomorphic registration of the two timepoints to create a midpoint average T1-image for each subject. Computing a mid-point average image prevents introduction of biases associated with using individual time-points as the reference (e.g., results differ depending of the time-point that is used as a reference). The mid-point average was scaled by the interscan interval. The registration step produced Jacobian determinant maps, which index contractions and expansions for each time-point relative to the mid-point average image. The default parameters for noise estimation, warping and bias regularizations were used. The mid-point average images were then segmented using SPM’s unified segmentation (Ashburner and Friston, 2005) with the following settings: light regularization, 60mm of FWHM gaussian smoothness of bias and 2,2,2,3,4,2 Gaussians per tissue type. The resulting 1.5mm^3^ GM and white matter maps were used to create separate DARTEL group templates for each sample (ADNI-1, and ADNI-GO/2). The DARTEL algorithm improves the inter-subject alignment by using multiple parameters per voxel when aligning the images (Ashburner, 2007). The segmented GM maps were multiplied by the Jacobian determinants of each time-point (Mechelli *et al.*, 2005). We then warped the segmented GM and white matter maps into the DARTEL template space with the ‘preserve amount’ option. Default settings were used elsewhere. The warping step produced modulated GM and white matter volumetric maps for each timepoint.

To check the quality of the images for subsequent analyses, the warped, modulated GM maps of each timepoint were visually inspected for segmentation and normalization inaccuracies by creating a plot showing a slice for all the subjects and were inspected for the presence of outliers by calculating the correlation between all the volumes using the CAT12 toolbox (http://dbm.neuro.uni-jena.de/cat/). Subjects that exhibited a correlation below 2 standard deviations and abnormalities on their segmented GM volume (e.g. low signal intensity, inhomogeneities, warping errors) were excluded. Two subjects were discarded due to low data quality in ADNI-1 and eight in ADNI-GO/2. See Table 2 for complete information of the included participants.

**Table 2.** Participant Demographics, APOE Genotype and CSF assays

| Profile | Dataset |  |  |  |  |  |
| --- | --- | --- | --- | --- | --- | --- |
|  | ADNI-1 |  |  | ADNI-GO/2 |  |  |
|  | aCSF | nCSF | Test | aCSF | nCSF | Test |
| *N* (Total) | 174 | 110 | $\chi^2=14.42^{***}$ | 316 | 237 | $\chi^2=11.28^{***}$ |
| CN/MCI/AD | 28/89/57 | 64/41/5 | $\chi^2=64.26^{***}$ | 39/130/147 | 162/75/0 | $\chi^2=230.84^{***}$ |
| Age | 74.51 (7.04) | 75.53 (6.5) | $t=-1.24$ | 72.16 (7.22) | 70.44 (6.56) | $t=2.91^{**}$ |
| Female/Male | 66/108 | 50/60 | $\chi^2=1.28$ | 146/170 | 110/127 | $\chi^2=0.00$ |
| Education | 15.57 (2.99) | 15.88 (2.89) | $t=-0.87$ | 15.96 (2.66) | 16.62 (2.56) | $t=2.96^{**}$ |
| $\epsilon 4+/\epsilon 4-$ | 124/50 | 16/94 | $\chi^2=84.48^{***}$ | 222/94 | 60/177 | $\chi^2=107.54^{***}$ |
| Aβ | 579.28 (179.15) | 1468.76 (554.28) | $t=-16.30^{***}$ | 641.66 (192.73) | 1376.54 (589.78) | $t=-18.45^{***}$ |
| pTau | 34.33 (12.33) | 19.6 (6.45) | $t=13.16^{***}$ | 35.75 (14.98) | 18.23 (6.52) | $t=18.57^{***}$ |

Information of the included participants from the ADNI-1 and ADNI-GO/2 cohorts by CSF group. Values represent means (standard deviations) unless otherwise specified. aCSF = abnormal CSF group; nCSF = normal CSF group; Aβ = CSF concentration of the amyloid beta 1-42 peptide; pTau = CSF concentration of hyper phosphorylated tau (at threonine 181). Independent samples t-tests (t) and chi-square ( $\chi^2$ ) tests were used to test CSF group differences within each ADNI Cohort. Baseline clinical diagnosis: CN=cognitively normal, MCI=mild cognitively impaired, AD=Alzheimer's disease. APOE status: $\epsilon 4+$ = $\epsilon 4$ carrier, $\epsilon 4-$ = $\epsilon 4$ non-carrier. Age and Education are in years. Aβ 1-42 and pTau concentrations are in picograms per milliliter (pg/mL). \*\*\* $p<0.001$ , \*\* $p<0.005$ .

### ROI definition

To test the predictive pathological models between the NbM and EC, we extracted GM volumes from two regions of interest (ROIs) using probabilistic atlases. Bilateral anatomical ROIs were created in MNI space with the SPM Anatomy Toolbox r 2.2b (http://www.fz-juelich.de/inm/inm-1/DE/Forschung/_docs/SPMAnatomyToolbox/SPMAnatomyToolbox_node.html) using previously published probabilistic maps of the NbM (Zaborszky *et al.*, 2008) and the EC (Amunts *et al.*, 2005). The NbM ROI is equivalent to areas Ch4 and Ch4p (Mesulam, 1983; Mesulam and Geula, 1988). The ROIs were then resampled and warped to the DARTEL group template (see Fig. 1b). Estimates of modulated GM volume (in units of milliliters) at each time-point were obtained using the ROI as a masking region, and by summing the total intensity of voxels falling within the masking region adjusted for the size and number of voxels (http://www0.cs.ucl.ac.uk/staff/g.ridgway/vbm/get_totals.m). No spatial smoothing was applied to the volumes. In addition, the total gray matter (TGM) from the modulated and warped individual time points was calculated using the same procedure. The total intracranial volume (TICV) was computed by summing up the tissue volumes of GM, white matter and CSF of the segmented average images produced in a previous step using the *seg8.mat file from the segmentation (Malone *et al.*, 2015).

In order to assess longitudinal changes in GM volumes, annualized percent change (APC) from baseline was calculated with the following formula (Cavedo *et al.*, 2017):

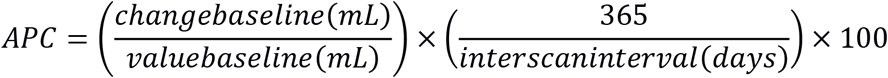

### Statistical Analyses

#### ANOVA

To compare baseline volumes and longitudinal volume changes between the CSF groups, we computed fixed effects ANOVAs for each ADNI cohort and each ROI with CSF Group (aCSF, nCSF) as a factor covarying for Age, Sex, Education, TICV, baseline TGM and APOE genotype (see Supplementary Fig. 1 for the bivariate correlation values among the modeled variables). We then pooled the data from both ADNI Cohorts and computed one way fixed effects ANOVAs with CSF group (aCSF, nCSF) as a factor, correcting for the same covariates, and an additional covariate for study Cohort (Ma *et al.*, 2018). All ANOVA models were carried out in R with a type III sums of squares.

#### Robust regression

To test the predictive staging models (NbM→EC and EC→NbM), robust linear regression analyses were carried out in Matlab R2018b with *filtm* and a bisquare weight function to minimize the influence of outliers. Each model covaried for Age, Sex, Education, TICV, baseline TGM and APOE genotype. The predictive models were calculated first for each dataset separately and then in the pooled sample collapsing across both Cohorts and including and additional covariate for Cohort.

#### Moderation analyses of independent samples

The moderation analyses (Fig. 3e and 3j) were carried out using the PROCESS macro for SPSS (Hayes, 2012) retrieved from http://www.afhayes.com/public/process2012.pdf.2012). The aCSF and nCSF groups served as the dichotomous moderator variable in all models. Age, Sex, Education, TICV, baseline TGM, APOE genotype and Cohort were included as covariates in all models. All data were mean centered. The moderation analysis employed a heteroscedasticity consistent standard error and covariance matrix estimator.

#### Comparison of correlations between dependent samples

To compare correlation coefficients between the NbM→EC and EC→NbM models, we used *cocor* to account for the dependence between the two models (Diedenhofen and Musch, 2015).

### Whole-brain analysis

To investigate the anatomical specificity of the relationship between baseline NbM volumes and longitudinal degeneration in the EC, we pursued an exploratory whole brain voxel-wise analysis. Whole-brain APC maps were calculated with the previous APC formula with the *imcalc* tool in SPM12. The APC maps were smoothed with a 4mm FWHM Gaussian kernel (Grothe *et al.*, 2012; Cantero *et al.*, 2017). For exploratory purposes, we tested a model with baseline EC predicting APC in every voxel in the aCSF group pooled across the ADNI-1 and ADNI-GO/ADNI-2 datasets. These regression models were estimated with non-parametric permutation testing using the Statistical NonParametric Mapping toolbox v13 (http://warwick.ac.uk/snpm) for SPM12 (Nichols and Holmes, 2002). Age, Sex, Education, TICV, baseline TGM, APOE genotype and Cohort were included as covariates. The models were estimated with 5000 permutations, without variance smoothing and global normalization, and including an explicit mask of the binarized and thresholded (>0.1) DARTEL GM ADNI-GO/2 group template. Results are presented with a voxel-level threshold of *p*<0.05 Family-Wise Error (FWE) corrected for multiple comparisons across the brain.

### Data availability

The data that support the findings of this study are openly available for request in LONI at https://ida.loni.usc.edu/login.jsp

## Results

### Abnormal CSF pTau/Aβ differentiates MRI indices of NbM and EC neurodegeneration

We first examined how the CSF pTau/Aβ grouping criteria differentiates patterns of baseline NbM and EC volume. Consistent with the sensitivity of CSF to neuropathology, we found that individuals with abnormal CSF cut-offs had smaller baseline volumes in the NbM in the ADNI-1 (main effect of CSF Group: *F*_1,276_=9.04, *p=*0.002), ADNI-GO/2 (main effect of CSF Group: *F*_*1,594*_=78.16, *p*<0.001) and in the pooled data of both cohorts (main effect of CSF Group: *F*_*1,828*_=75.88, *p*<0.001; Fig 2a). Baseline volumes of the EC were also smaller in the aCSF group in ADNI-1 (main effect of CSF Group: *F*_*1,276*_=8.71, *p=*0.003), ADNI-GO/2 (main effect of CSF Group: *F*_*1,594*_=21.98, *p*<0.001) and in the pooled data (main effect of CSF Group: *F*_*1,828*_=25.86, *p*<0.001; Fig 2b).

**Figure 2.**
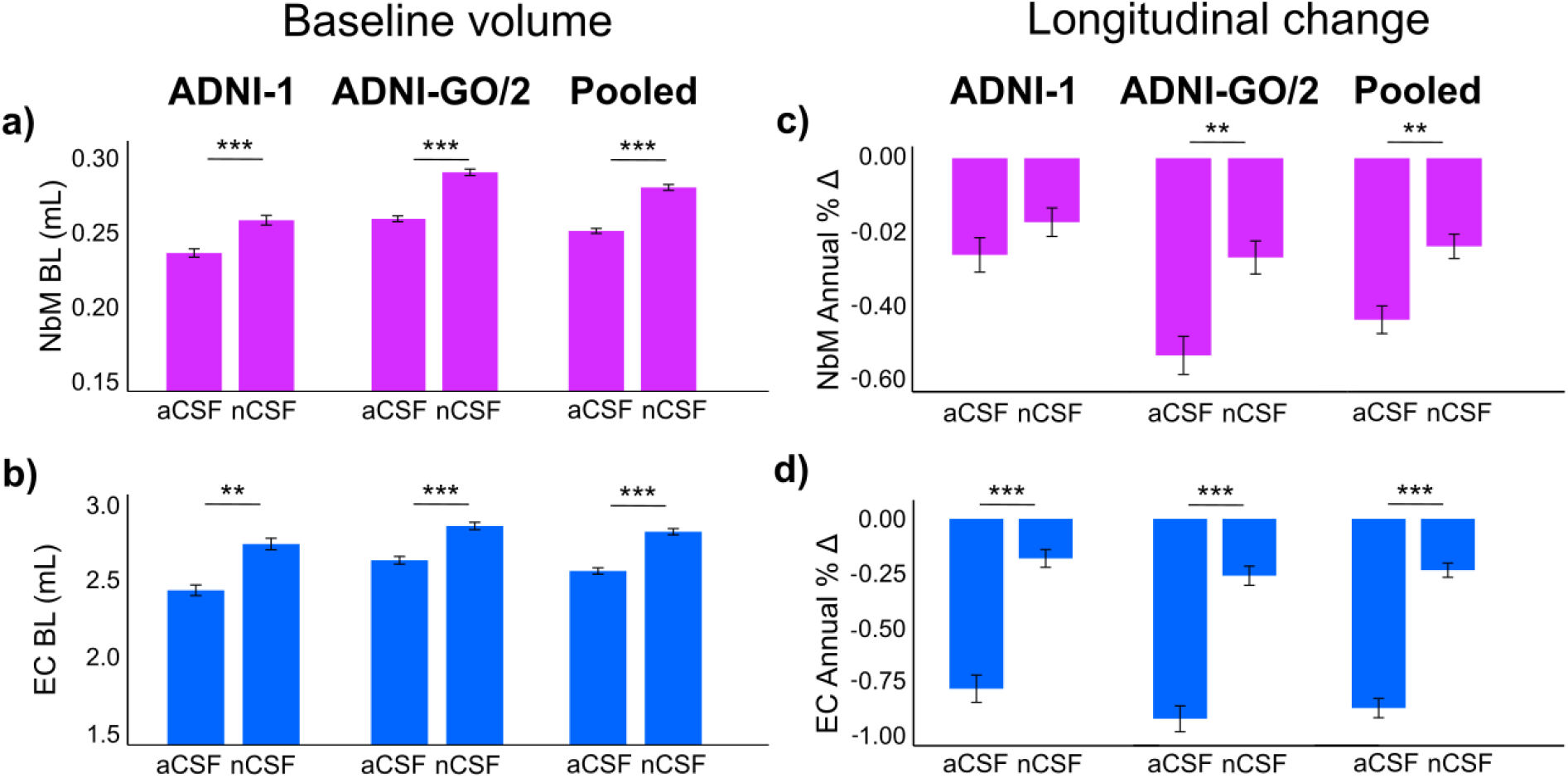
CSF biomarkers of neuropathology differentiate NbM and EC indices of structural degeneration. Bar plots showing the means and standard errors of the baseline and longitudinal changes of the NbM (purple bars) and EC (blue bars) a priori ROIs in the two independent datasets (ADNI-1 and ADNI-GO/2) and the pooled data of both datasets. Baseline NbM (a) and EC (b) volumes were smaller in the aCSF group. Longitudinal degeneration in the NbM (c) and in the EC (d) was larger in the aCSF group. aCSF=abnormal CSF group; nCSF=normal CSF group; NbM=nucleus basalis of Meynert, EC=entorhinal cortex. mL=milliliters, BL=baseline volume, asterisks denote statistical significance (\*\*\**p*<0.001; \*\**p*<0.01; \**p*<0.05).

We next examined if the CSF pTau/Aβ grouping criteria influenced longitudinal indices of APC in the NbM and EC at each ADNI Cohort. These models revealed that individuals with abnormal CSF biomarkers showed more longitudinal atrophy in the NbM in the ADNI-GO/2 (main effect of CSF Group: *F*_*1,594*_=7.60, *p=*0.005) and in the pooled dataset (main effect of CSF Group: *F*_*1,828*_=9.79, *p=*0.002), but not in ADNI-1 (main effect of CSF Group: *F*_*1,276*_=2.54, *p=*0.11; Fig 2c). In the EC, the aCSF group exhibited more longitudinal atrophy in ADNI-1 (main effect of CSF Group: *F*_*1,276*_=23.91, *p<*0.001), ADNI-GO/2 (main effect of CSF Group: *F*_*1,594*_=29.54, *p<*0.001) and in the pooled data (main effect of CSF Group: *F*_*1,828*_=47.51, *p<*0.001; Fig 2d). Finally, we investigated if the magnitude of longitudinal degeneration in the NbM and in the EC differed between individuals at different disease stages. For that, we cross-referenced the CSF groups with the ADNI baseline Clinical Diagnosis and found that, individuals with abnormal CSF biomarkers in all disease stages had increased degeneration in the NbM and the EC. Crucially, individuals in the preclinical disease stage (aCSF and normal cognition at baseline) showed greater magnitudes of degeneration in the NbM and the EC compared to the cognitively normal controls (nCSF and normal cognition at baseline). See Supplementary Fig. 2 and Supplementary Table 1. Consistent with our hypotheses, CSF biomarkers of abnormal pTau/Aβ accumulation in the central nervous system differentiated both baseline gray matter volume and longitudinal gray matter degeneration in the NbM and EC.

### Abnormal pTau/AB potentiates the predictive spread of NbM→EC degeneration

Thus far, our multimodal CSF and structural MRI analyses have treated patterns of neurodegeneration in the NbM and EC separately. We next used regression-based modelling to test if the degeneration in the NbM and in the EC reflects interdependent pathological processes. If degeneration in the NbM precedes degeneration in the EC due to selective neuronal vulnerability of cholinergic neurons inducing a ‘spread’ of pathology, then NbM baseline volumes should predict degeneration in the EC (NbM→EC). However, if degeneration in NbM and EC occurs in parallel due to a common global pathology, then baseline volumes of either region should predict degeneration in the other. Alternatively, if degeneration in NbM and EC reflects independent local pathology, then baseline volumes of neither region should predict degeneration in the other.

We used robust linear regression to examine these competing models in the aCSF and nCSF groups. Consistent with an interdependent NbM→EC model, we found in the aCSF group that baseline NbM volumes predicted longitudinal degeneration in the EC in both ADNI-1 (Fig. 3a; r=0.20, t_166_=2.12, p=0.03) and ADNI-GO/2 (Fig. 3b; r=0.37, t_308_=5.15, p<0.001). We next examined whether the predictive pathological staging of NbM and EC degeneration is observable under conditions of normal CSF pTau/Aβ. It could be the case that the underlying neuropathological process driving this relationship also occurs at ‘neurotypical’ levels of pTau/Aβ, but that this process is exacerbated in the aCSF group. In this case, we would expect to observe the same NbM→EC interrelationship, though possibly at a smaller magnitude. We found negligible evidence for the NbM→EC model in the nCSF group in ADNI-1 (Fig.3c; r=-0.17, t_102_=-1.14, p=0.25). However, we did observe a predictive relationship in ADNI-GO/2 (Fig.3d, r=0.21, t_229_=2.47, p=0.01). We then examined these relationships in the pooled data. The results supported the NbM→EC model in the aCSF (r=0.30, t_481_= 5.32, p<0.001) but not the nCSF group (r = 0.11, t_338_=1.45, p=0.14). See Figure 3e and Supplementary Table 2. To exclude the possibility that the results were distorted by possible collinearities among the model predictors and covariates, we computed each model without any of the covariates. The exclusion of covariates did not alter any of the above findings (see Supplementary Table 3). To directly test if the relationship observed in the pooled aCSF group was significantly different from the that observed in the pooled nCSF group, we computed a moderation analysis with CSF group (aCSF, nCSF) as a dichotomous moderator on the NbM→EC interrelationship (see Methods). We found a significant moderating effect of CSF group on the NbM→EC interrelationship (*t*_826_=2.55, *p*=0.01), indicating a more pronounced interregional spread in the aCSF compared to the nCSF group.

**Figure 3.**
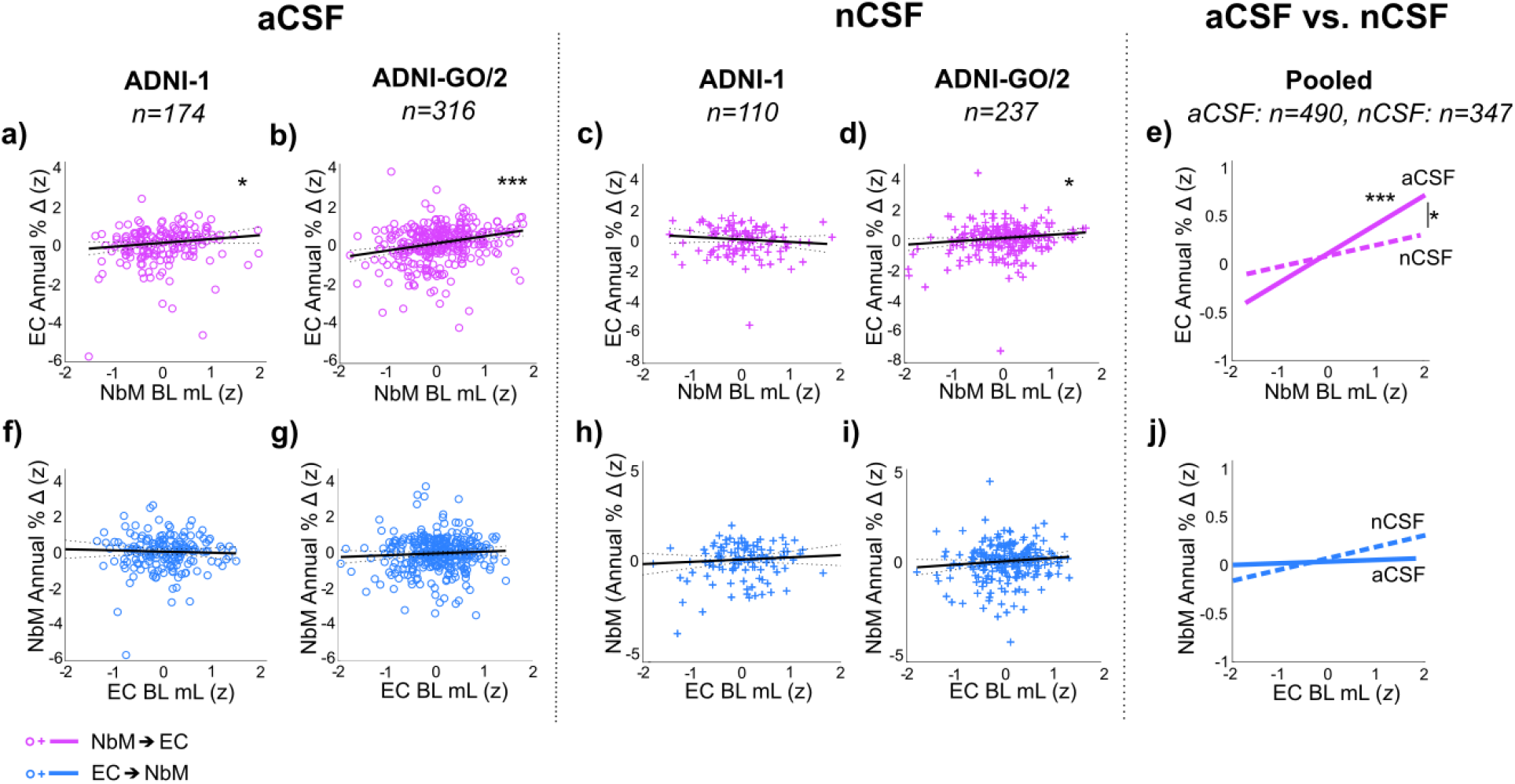
Predictive pathological staging models. In the aCSF group, smaller baseline NbM volumes (x-axis: negative z-score values) were associated with greater magnitudes of longitudinal degeneration in the EC (y-axis: negative z-score values for T2-T1 annual percent change) in both (a) ADNI-1 and (b) ADNI-GO/2 (lower left corner of the graphs). No such relationships were observed in the nCSF groups in (c) ADNI-1 but it was present in (d) ADNI-GO/2. (e) CSF concentrations of pTau/Aβ (nCSF versus aCSF) significantly moderated the relationship between baseline NbM volume and longitudinal EC degeneration. (f—j) Baseline EC volumes did not predict longitudinal degeneration in the NbM in either the aCSF or nCSF, in either ADNI-1 or ADNI-GO/2. Nor did it moderate this relationship. aCSF=abnormal CSF group; nCSF=normal CSF group; NbM=nucleus basalis of Meynert, EC=entorhinal cortex. mL=milliliters, BL=baseline volume, asterisks denote statistical significance (\*\*\**p*<0.001;\**p*<0.01; * *p*<0.05).

We next tested the competing model of EC→NbM predictive pathological spread, focusing first on the aCSF group. We found that EC baseline volumes did not predict longitudinal decreases in the NbM in ADNI-1 (Fig. 3f; r=-0.06, t_166_=-0.56, p=0.57), in ADNI-GO/2 (Fig. 3g; r=0.10, t_308_=1.18, p=0.23) or in the pooled sample (Fig.3j; r=0.02, t_481_=0.26, p=0.79; Supplementary Table 1). Moreover, we found negligible evidence for the EC→NbM model in the nCSF group in ADNI-1 (Fig.3h; r=0.12, t_102_=0.92, p=0.36), ADNI-GO/2 (Fig.3i, r=0.18, t_229_=1.72, p=0.08), or in the pooled sample (Fig.3j; r=0.12, t_334_=1.53, p=0.12; Supplementary Table 1). Nor was there a significant interaction between CSF group and the relationship between EC and NbM in the pooled sample (Fig 3j; *t*_826=_1, *p*=0.62).

We then directly compared the two predictive staging models, NbM→EC and EC→NbM, using a slopes analysis between dependent samples (see Methods). We found that the coefficients in each model were significantly different from one another in the aCSF group (*z* = 5.54; *p*<0.001; 95% r CI: 0.1831-0.3847) but not in the nCSF group (*z* = 0.08; *p=*0.931; 95% r CI: −0.1424 −0.13). See Figure 4.

**Figure 4.**
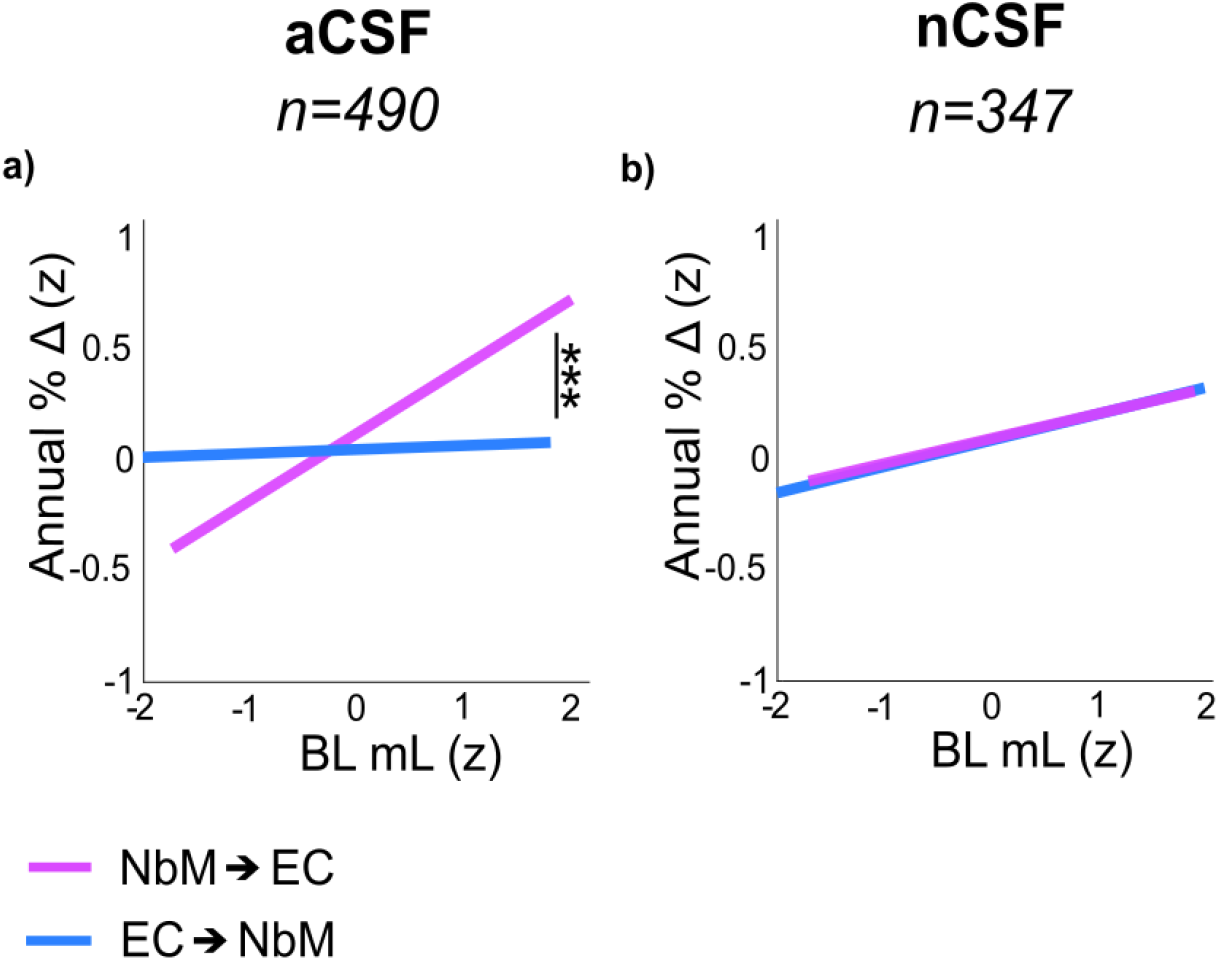
Aβ and pTau potentiate the spread of neurodegeneration from NbM to EC. The slopes of the NbM→EC and EC→NbM robust regression models were different in (a) the pooled aCSF group but (b) not in the pooled nCSF group. The x axes reflect baseline NbM volumes (purple lines) and baseline EC volumes (blue lines). The y axes reflect annual percent change in the EC (purple lines) and in the NbM (blue lines). Baseline volumes and annual percent change units were z-scored (z). aCSF=abnormal CSF group; nCSF=normal CSF group; NbM=nucleus basalis of Meynert, EC=entorhinal cortex. mL=milliliters, BL=baseline volume, asterisks denote statistical significance (\*\*\**p*<0.001;\**p*<0.01; * *p*<0.05).

Taken together, our predictive pathological models suggest that neurodegeneration of the cholinergic basal forebrain system may occur in both the aCSF and nCSF aging phenotypes, but that abnormal proteopathic pTau/Aβ accumulation greatly exacerbates this effect. The results also demonstrate that the EC→NbM model does not fit a neurobiologically plausible predictive staging of neurodegeneration in either the aCSF or nCSF cohort.

### Clinical diagnosis does not moderate the relationship between NbM→EC

Having demonstrated that the NbM→EC relationship was more prominent and reliable in the aCSF group, we then investigated whether the severity of the disease had an impact on the strength of this relationship in the pooled aCSF group. We expected that in initial (i.e. cognitively normal aCSF individuals) and advanced disease stages (i.e. cognitively impaired aCSF individuals), participants would show a weaker predictive relationship due to smaller magnitudes of EC atrophy in the early stages and reduced NbM atrophy in the more advanced stages (Schmitz and Spreng, 2016). Clinical diagnosis was not a significant moderating effect (Supplementary Figure 3; *t*_*1,479*_=1.65, p=0.49), indicating that the predictive relationship was robust across clinical diagnostic subgroups of the aCSF cohort.

### NbM→EC predictive pathological spreading is anatomically specific

How spatially selective is the observed interrelationship between NbM and EC in the aCSF group? To determine where degeneration is predicted based on NbM integrity, we performed a voxel-wise regression analysis between baseline volumes in the NbM and longitudinal APC in every brain voxel (see Methods). The model collapsed aCSF individuals across the two ADNI Cohorts into a single cohort. We found that baseline NbM volumes predicted degeneration in a circumscribed cluster of regions localized to the right entorhinal and perirhinal cortices and in the left middle temporal gyrus (Fig. 5a). The predictive pathological spread of NbM→EC neurodegeneration therefore exhibits a high degree of anatomical specificity.

**Figure 5.**
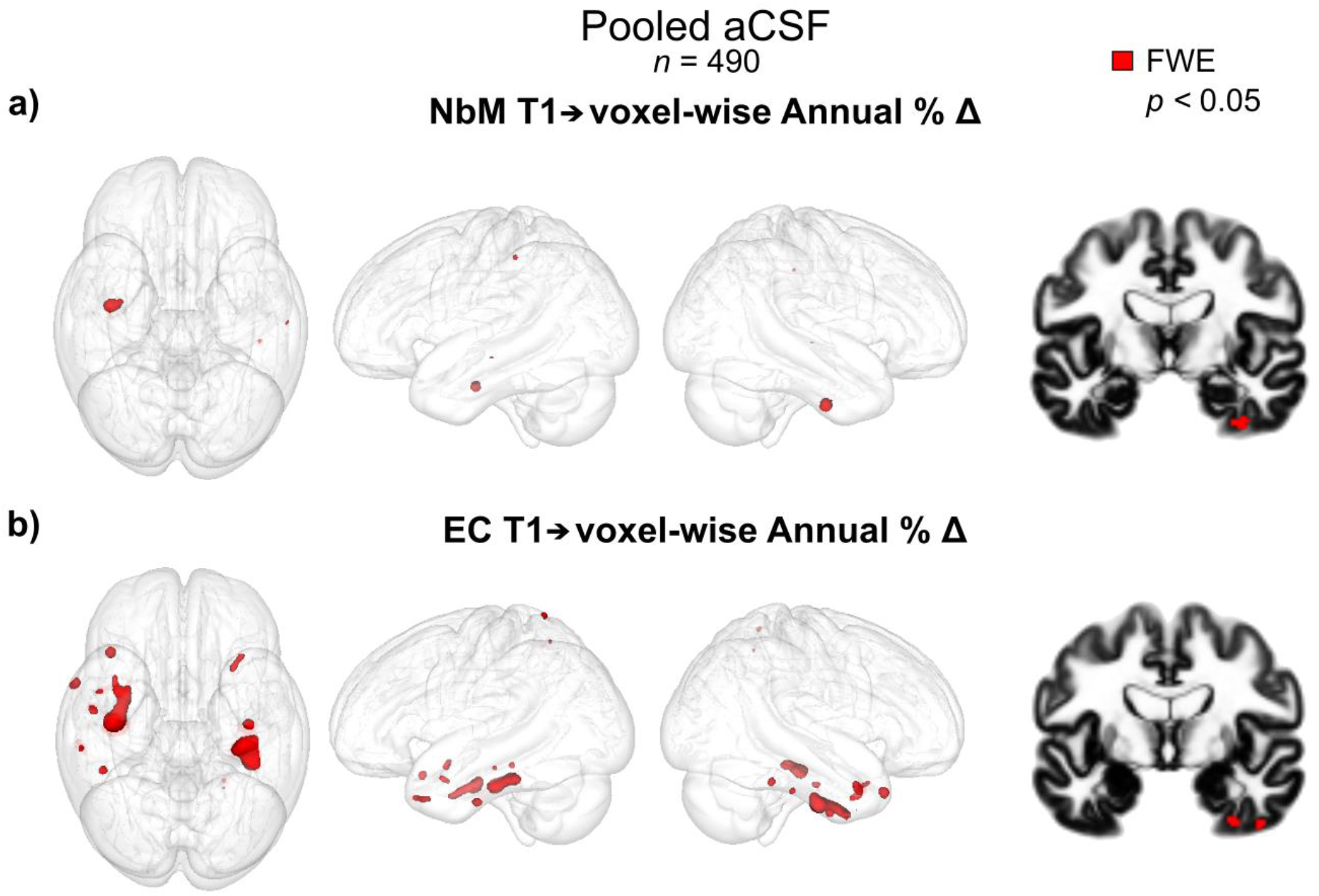
NbM and EC predict distinct patterns of downstream degenerative spreading. Baseline volumes of the NbM and the EC predicting APC in every brain voxel in the pooled aCSF group (n=490). In red, voxels showing a significant association with baseline volumes in the NbM (a) and in the EC (b) with a *p*<0.05 voxel-level FWE-corrected threshold. Baseline NbM volumes predicted degeneration in the EC whereas baseline EC volumes predicted degeneration in regions of the temporal and parietal cortices. Results are rendered into an MNI glass brain to the group DARTEL space using Mango (http://ric.uthscsa.edu/mango/). Models were tested for significance using non-parametric permutation testing. aCSF=abnormal CSF group; NbM=nucleus basalis of Meynert, EC=entorhinal cortex; FWE=Family-Wise Error.

In our a priori ROI-based regression models, we found negligible evidence in support of the EC→NbM model. However, following from non-human animal work (Khan *et al.*, 2014), we hypothesized that that predictive pathological spread of degeneration from EC might target anatomically and functionally connected cortical areas including the parahippocampal and posterior parietal cortices. To test this hypothesis, we computed a second voxel-wise regression analysis in the pooled aCSF group between baseline volumes in the EC and longitudinal APC in every brain voxel of the gray matter mask (see Methods). We found that baseline EC volumes predicted longitudinal degeneration in a circumscribed network of temporal cortical regions including the entorhinal, perirhinal, parahippocampal and fusiform cortices and temporal poles (Figure. 5b). Additional clusters were detected in the insular and posterior parietal cortices.

Finally, we tested the same models in the nCSF cohort, collapsing individuals from both ADNI Cohorts. No significant voxels were found at the corrected statistical threshold (*p<*0.05 FWE) in either of the models (NbM→whole brain; EC→whole brain). Taken together, the voxel-wise regression results show that among individuals expressing abnormal CSF: (1) baseline NbM volumes selectively predict degeneration in the EC and adjacent perirhinal cortex, and, (2) EC degeneration ‘spreads’ to anatomically and functionally connected areas including the parahippocampal gyrus and other temporoparietal regions (Khan *et al.*, 2014).

## Discussion

We provide evidence that degeneration in the NbM is an upstream event of subsequent degeneration in the EC and temporal cortices. Our results generalized across two non-overlapping and well-powered datasets. As hypothesized, the NbM→EC relationship was moderated by the presence of abnormal CSF biomarkers of pTau and Aβ, consistent with a mechanism of trans-synaptic spread of these proteins across anatomically connected regions. Using whole-brain regression models, we presented novel evidence that the relationship between baseline NbM volumes and neurodegeneration was spatially specific to regions of the EC and the perirhinal cortices, regions hypothesized to be among the earliest to be affected in Alzheimer’s disease (Braak and Braak, 1991; Khan *et al.*, 2014). Finally, baseline EC volumes predicted neurodegeneration mainly in regions of the temporal cortex, recapitulating the prevailing staging model (Braak and Braak, 1991; Braak *et al.*, 2006; Liu *et al.*, 2012). These findings highlight the relevance of the cholinergic BF nuclei in Alzheimer’s disease and motivate novel interventions targeted to these neuronal populations.

The intrinsic properties of the BF cholinergic neurons, including their location, connections and morphology, might be key to understanding why these cell types are selectively vulnerable in Alzheimer’s disease (Mattson and Magnus, 2006). The NbM cholinergic neurons have large projecting axons with extensive arbors that expand long distances in the central nervous system (Wu *et al.*, 2014; Ballinger *et al.*, 2016). Extremely large axons and arbors exert high metabolic demands in order to support their maintenance, repair and transport. A less efficient cellular response to axonal perturbations could increase their vulnerability to the accumulation of abnormal proteins (Wu *et al.*, 2014). In addition, large surface areas increase their exposure to toxic environments (Mattson and Magnus, 2006). Consistent with its selective vulnerability, the NbM is one of the most vulnerable regions to the early accumulation of intracellular tau pathology (Mesulam *et al.*, 2004; Braak and Del Tredici, 2011) and, interestingly, intraneuronal amyloid oligomers (Baker-Nigh *et al.*, 2015). There is increasing evidence showing that intraneuronal amyloid can have deleterious early effects in the cells, such as disrupting their retrograde axonal transport machinery (Morfini *et al.*, 2009) which in turn can cause a neurotrophic deficiency (Bellucci *et al.*, 2006) leading to neurodegeneration (De Lacalle *et al.*, 1996). *In vivo* structural MRI studies have added further support for the vulnerability of the BF, showing reduced BF volumes cross-sectionally (Grothe *et al.*, 2012; Cantero *et al.*, 2017) and longitudinally (Hall *et al.*, 2008; Grothe *et al.*, 2013; Schmitz *et al.*, 2018) in individuals in the Alzheimer’s disease continuum.

In this study, the pathological spread of neurodegeneration from the NbM to the EC was exacerbated in individuals with abnormal levels of CSF pTau and Aβ. Evidence from non-human animal studies shows that protein aggregates can spread trans-synaptically to neuroanatomically connected regions. In an amyloid precursor protein mouse model of Alzheimer’s disease, neuronal accumulation of Aβ was observed first in the EC, followed by the dentate gyrus, a downstream region receiving monosynaptic projections from the EC (Harris *et al.*, 2010). Evidence further suggests that tau aggregates propagate from the EC to limbic and then neocortical connected areas (De Calignon *et al.*, 2012; Kfoury *et al.*, 2012; Walker *et al.*, 2013; Wu *et al.*, 2016). For example, in mice that expressed human tau restricted to the EC, the axonal terminals of the EC projecting neurons showed increasing markers of tauopathy over time (Liu *et al.*, 2012). In all, this shows that Aβ and tau can propagate synaptically across neural networks of anatomically connected vulnerable cells. If the NbM is one of the earliest regions to accumulate tau and Aβ, then these proteins may spread to the EC trans-synaptically, inducing the observed downstream neurodegeneration. Our results are therefore compatible with animal research showing a trans-synaptic mechanism of abnormal spreading of tau proteins. However, to ultimately demonstrate that degeneration in the EC is driven by the spread of tau pathology from the NbM in humans, longitudinal studies integrating both structural MRI and tau positron emission tomography are needed. In addition, we observed a relationship between the NbM→EC in the normal CSF group. This effect was weaker and not reliable across samples, indicating the possibility that there are other age-related pressures on the NbM cholinergic neurons independent from Aβ and pTau (Nitsch *et al.*, 1992; Wurtman, 1992).

Using longitudinal whole-brain structural MRI, we also showed a reliable (Figure 3A—E) and highly specific (Figure 5A) sequence of neurodegeneration from the NbM to the EC. This result challenges the most widely accepted model of Alzheimer’s disease staging, which posits that the neuropathological and neurodegenerative cascade initiates in the transentorhinal and EC (Braak and Braak, 1991; Corder *et al.*, 2000; Braak *et al.*, 2006; Liu *et al.*, 2012; Khan *et al.*, 2014). This model was primarily advanced from *post-mortem* histology studies that examined the brains of individuals who died at different ages from different causes, which showed tau aggregates forming in the EC even in non-demented individuals (Braak and Braak, 1991; Braak *et al.*, 2006). Predictive pathological staging based on histological strategies are limited by the heterogeneity of the post-mortem samples, the inability to track pathology with progressive neurodegeneration within individuals, the variability in tissue staining techniques used to profile underlying pathology, and the variability in anatomical coverage across both cortical and subcortical regions. We overcame these limitations by using an *in vivo* longitudinal neuroimaging strategy of predictive pathological staging in a large cohort of over 800 older adults. We quantified the baseline CSF profile of amyloid and tau pathology in each individual based on the automated Elecsys protocol, which was harmonized between samples. We then applied an independently defined cutpoint to delineate unbiased groups of individuals exhibiting abnormal and normal CSF biomarkers. By comparing the spreading dynamics of neurodegeneration between these two groups, we demonstrated that baseline NbM volumes predicted an accelerated annual rate of degeneration in a future time point in the EC in the presence of abnormally elevated Aβ and pTau. In addition, we found evidence that this neuropathological background was associated with neurodegenerative spreading from EC to targets of its projections in the medial temporal lobes. These latter findings parallel those of recent cross-species translational research comparing transgenic mouse models of Alzheimer’s disease with at-risk human participants (Khan *et al.*, 2014). In all, our results added evidence that NbM degeneration is an earlier pathological event in the Alzheimer’s disease pathological cascade.

Here, we showed that the NbM→EC model reliably generalized between two different populations. It is important to note that the data of the ADNI-GO/2 sample were acquired in 3T and the ADNI-1 in 1.5T scanners. Previous reports have only used 1.5 datasets (Schmitz and Spreng, 2016) and it is well known that the signal to noise ratio of subcortical structures is higher at 3T (Tardif *et al.*, 2010; Marchewka *et al.*, 2014). It is therefore likely that degeneration in the NbM is more accurately quantified in the ADNI-GO/2 sample. Although it is estimated that approximately 90% of the NbM neurons in monkeys are cholinergic (Mesulam, 1983), to ultimately prove that the degeneration in the EC is driven by a cholinergic deficit in humans, degeneration that is specific to cholinergic cell types should be demonstrated. The combination of structural MRI with cholinergic positron emission tomography (Aghourian *et al.*, 2017; Schmitz *et al.*, 2018) could help elucidate this issue. Finally, late onset Alzheimer’s disease is a complex, multifactorial (Veitch *et al.*, 2018), polygenic (Tan *et al.*, 2019) neurodegenerative disease. We found that the relationship between NbM→EC was robust to differences in APOE genotype and clinical diagnosis—the presence of Aβ and pTau was the most potent moderating factor observed in our study. Nevertheless, other inter-individual variables not modeled in the present study such as vascular health (Iturria-Medina *et al.*, 2016) could help gain additional insights on the selective vulnerability of the NbM and the EC to Alzheimer’s disease.

In conclusion, the current study provides reliable and anatomically specific support for a predictive pathological staging model of Alzheimer’s disease that progresses in the following sequence: NbM→EC→Neocortex. In all, our findings call into question the prevailing view of Alzheimer’s disease pathogenesis, suggesting instead that degeneration of the basal forebrain cholinergic projection system is an upstream event to that of its most vulnerable cortical targets in the EC. If the cholinergic neurons of the BF are selectively vulnerable to the early accumulation and propagation of Alzheimer’s disease pathology, then interventions targeting these neuronal populations in early disease stages may be useful for modifying disease progression prior to cognitive decline.

## Abbreviations

aCSF: Abnormal CSF group based on the pTau/Aβ_1-42_ ratio >= 0.28
ADN: Alzheimer’s Disease Neuroimaging Initiative
APC: Annual Percent Change
Aβ: Amyloid Beta
BF: Basal forebrain
EC: Entorhinal Cortex
GM: Gray Matter
NbM: Nucleus Basalis of Meynert
nCSF: Normal CSF group based on the pTau/Aβ_1-42_ ratio <0.28
pTau: Hyper phosphorylated tau
sMRI: Structural MRI
ROI: Region of Interest
TGM: Total Gray Matter
TICV: Total Intracranial Volume

## Acknowledgements

Data collection and sharing for this project was funded by the Alzheimer’s Disease Neuroimaging Initiative (ADNI) (National Institutes of Health Grant U01 AG024904) and DOD ADNI (Department of Defense award number W81XWH-12-2-0012). ADNI is funded by the National Institute on Aging, the National Institute of Biomedical Imaging and Bioengineering, and through generous contributions from the following: AbbVie, Alzheimer’s Association; Alzheimer’s Drug Discovery Foundation; Araclon Biotech; BioClinica, Inc.; Biogen; Bristol-Myers Squibb Company; CereSpir, Inc.; Cogstate; Eisai Inc.; Elan Pharmaceuticals, Inc.; Eli Lilly and Company; EuroImmun; F. Hoffmann-La Roche Ltd and its affiliated company Genentech, Inc.; Fujirebio; GE Healthcare; IXICO Ltd.; Janssen Alzheimer Immunotherapy Research & Development, LLC.; Johnson & Johnson Pharmaceutical Research & Development LLC.; Lumosity; Lundbeck; Merck & Co., Inc.; Meso Scale Diagnostics, LLC.; NeuroRx Research; Neurotrack Technologies; Novartis Pharmaceuticals Corporation; Pfizer Inc.; Piramal Imaging; Servier; Takeda Pharmaceutical Company; and Transition Therapeutics. The Canadian Institutes of Health Research is providing funds to support ADNI clinical sites in Canada. Private sector contributions are facilitated by the Foundation for the National Institutes of Health (www.fnih.org). The grantee organization is the Northern California Institute for Research and Education, and the study is coordinated by the Alzheimer’s Therapeutic Research Institute at the University of Southern California. ADNI data are disseminated by the Laboratory for Neuro Imaging at the University of Southern California. This work was supported in part by a grant from the NIA (R03 RAG060263A) (T.W.S. and R.N.S.) and the Canada First Research Excellence Fund (T.W.S. and R.N.S.). S.F.C was supported by the Doctoral College “Imaging the Mind” (FWF-W1233) of the Austrian Science Fund. We thank Didac Vidal-Piñeiro and the LCBC group in Oslo for feedback on this manuscript.

The authors declare no competing financial interests.

## References

Aghourian M, Legault-Denis C, Soucy JP, Rosa-Neto P, Gauthier S, Kostikov A, et al. Quantification of brain cholinergic denervation in Alzheimer’s disease using PET imaging with [18F]-FEOBV. Mol Psychiatry 2017; 22: 1531

Aisen PS, Petersen RC, Donohue M, Weiner MW. Alzheimer’s Disease Neuroimaging Initiative 2 Clinical Core: Progress and plans. Alzheimer’s Dement 2015; 11: 734–739.

Aisen PS, Petersen RC, Donohue MC, Gamst A, Raman R, Thomas RG, et al. Clinical core of the Alzheimer’s disease neuroimaging initiative: Progress and plans. Alzheimer’s Dement 2010; 6: 239–246.

Amunts K, Kedo O, Kindler M, Pieperhoff P, Mohlberg H, Shah NJ, et al. Cytoarchitectonic mapping of the human amygdala, hippocampal region and entorhinal cortex: Intersubject variability and probability maps. Anat Embryol (Berl) 2005; 210: 343–352.

Ashburner J. A fast diffeomorphic image registration algorithm. Neuroimage 2007; 38: 95–113.

Ashburner J, Friston KJ. Unified segmentation. Neuroimage 2005; 26: 839–851.

Ashburner J, Ridgway GR. Symmetric diffeomorphic modeling of longitudinal structural MRI. Front Neurosci 2013; 6: 1–19.

Baker-Nigh A, Vahedi S, Davis EG, Weintraub S, Bigio EH, Klein WL, et al. Neuronal amyloid-β accumulation within cholinergic basal forebrain in ageing and Alzheimer’s disease. Brain 2015; 138: 1722–1737.

Ballinger EC, Ananth M, Talmage DA, Role LW. Review Basal Forebrain Cholinergic Circuits and Signaling in Cognition and Cognitive Decline. Neuron 2016; 91: 1199–1218.

Bellucci A, Luccarini I, Scali C, Prosperi C, Giovannini MG, Pepeu G, et al. Cholinergic dysfunction, neuronal damage and axonal loss in TgCRND8 mice. Neurobiol Dis 2006; 23: 260–272

Braak H, Alafuzoff I, Arzberger T, Kretzschmar H, Tredici K. Staging of Alzheimer disease-associated neurofibrillary pathology using paraffin sections and immunocytochemistry. Acta Neuropathol 2006; 112: 389–404.

Braak H, Braak E. Neuropathological stageing of Alzheimer-related changes. Acta Neuropathol 1991; 82: 239–59.

Braak H, Del Tredici K. The pathological process underlying Alzheimer’s disease in individuals under thirty. Acta Neuropathol 2011; 121: 171–181.

De Calignon A, Polydoro M, Suárez-Calvet M, William C, Adamowicz DH, Kopeikina KJ, et al. Propagation of Tau Pathology in a Model of Early Alzheimer’s Disease. Neuron 2012; 73: 685–697

Cantero JL, Zaborszky L, Atienza M. Volume loss of the nucleus basalis of meynert is associated with atrophy of innervated regions in mild cognitive impairment. Cereb Cortex 2017; 27: 3881–3889.

Cavedo E, Grothe MJ, Colliot O, Lista S, Chupin M, Dormont Di, et al. Reduced basal forebrain atrophy progression in a randomized Donepezil trial in prodromal Alzheimer’s disease. Sci Rep 2017; 7: 1–10.

Corder EH, Saunders AM, Strittmatter WJ, Schmechel DE, Gaskell PC, Small GW, et al. Gene dose of apolipoprotein E type 4 allele and the risk of Alzheimer’s disease in late onset families. Science (80-) 1993; 261: 921–923.

Corder EH, Woodbury MA, Volkmann I, Madsen DK, Bogdanovic N, Winblad B. Density profiles of Alzheimer disease regional brain pathology for the Huddinge brain bank: Pattern recognition emulates and expands upon Braak staging. In: Experimental Gerontology. 2000. p. 851–864

Crane PK, Carle A, Gibbons LE, Insel P, Mackin RS, Gross A, et al. Development and assessment of a composite score for memory in the Alzheimer’s Disease Neuroimaging Initiative (ADNI). Brain Imaging Behav 2012; 6: 502–516.

Diedenhofen B, Musch J. Cocor: A comprehensive solution for the statistical comparison of correlations. PLoS One 2015; 10: 1–12.

Gibbons. A composite score for executive functioning, validated in Alzheimer’s Disease Neuroimaging Initiative (ADNI) participants with baseline mild cognitive impairment. Brain Imaging Behav 2012; 6: 517–527.

Grothe M, Heinsen H, Teipel S. Longitudinal measures of cholinergic forebrain atrophy in the transition from healthy aging to Alzheimer’s disease. Neurobiol Aging 2013; 34: 1210–1220.

Grothe M, Heinsen H, Teipel SJ. Atrophy of the cholinergic basal forebrain over the adult age range and in early stages of Alzheimer’s disease. Biol Psychiatry 2012; 71: 805–813.

Hall AM, Moore RY, Lopez OL, Kuller L, Becker JT. Basal forebrain atrophy is a presymptomatic marker for Alzheimer’s disease. Alzheimer’s Dement 2008; 4: 271–279.

Hansson O, Seibyl J, Stomrud E, Zetterberg H, Trojanowski JQ, Bittner T, et al. CSF biomarkers of Alzheimer’ s disease concord with amyloid-b PET and predict clinical progression : A study of fully automated immunoassays in BioFINDER and ADNI cohorts. Alzheimers Dement 2018; 14: 1470–1481

Harris JA, Devidze N, Verret L, Ho K, Halabisky B, Thwin MT, et al. Transsynaptic Progression of Amyloid-β-Induced Neuronal Dysfunction within the Entorhinal-Hippocampal Network. Neuron 2010; 68: 428–441.

Hayes AF. PROCESS: A versatile computational tool for observed variable mediation, moderation, and conditional process modeling. 2012.

Iturria-Medina Y, Sotero RC, Toussaint PJ, Mateos-Pérez JM, Evans AC, Weiner MW, et al. Early role of vascular dysregulation on late-onset Alzheimer’s disease based on multifactorial data-driven analysis. Nat Commun 2016; 7: 11934

Kfoury N, Holmes BB, Jiang H, Holtzman DM, Diamond MI. Trans-cellular propagation of Tau aggregation by fibrillar species. J Biol Chem 2012; 287: 19440–19451.

Khan UA, Liu L, Provenzano FA, Berman DE, Profaci CP, Sloan R, et al. Molecular drivers and cortical spread of lateral entorhinal cortex dysfunction in preclinical Alzheimer’s disease. Nat Neurosci 2014; 17: 304–311.

De Lacalle S, Cooper JD, Svendsen CN, Dunnett SB, Sofroniew M V. Reduced retrograde labelling with fluorescent tracer accompanies neuronal atrophy of basal forebrain cholinergic neurons in aged rats. Neuroscience 1996; 75: 19–27

Liu C, Bu G. Apolipoprotein E and Alzheimer disease : risk, mechanisms and therapy. Nat Rev Neurol 2013; 9: 106

Liu L, Drouet V, Wu JW, Witter MP, Small SA, Clelland C, et al. Trans-synaptic spread of tau pathology in vivo. PLoS One 2012; 7

Ma D, Popuri K, Bhalla M, Jacova C, Wang L, Beg MF, et al. Quantitative Assessment of Field Strength, Total Intracranial Volume, Sex and Age on The Goodness of Harmonization For Volumetric Analysis on The ADNI Database Conclusion. Hum Brain Mapp 2018; 40: 1507–1527

Malone IB, Leung KK, Clegg S, Barnes J, Whitwell JL, Ashburner J, et al. Accurate automatic estimation of total intracranial volume: A nuisance variable with less nuisance. Neuroimage 2015; 104: 366–372.

Marchewka A, Kherif F, Krueger G, Grabowska A, Frackowiak R, Draganski B. Influence of magnetic field strength and image registration strategy on voxel-based morphometry in a study of Alzheimer’s disease. Hum Brain Mapp 2014; 35: 1865–1874.

Mattson MP, Magnus T. Ageing and neuronal vulnerability. Nat Rev Neurosci 2006; 7: 278–294.

Mechelli A, Price CJ, Friston KJ, Ashburner J. Voxel-Based Morphometry of the Human Brain: Methods and Applications. Curr Med Imaging Rev 2005; 1: 115–113

Mesulam M -M, Geula C. Nucleus basalis (Ch4) and cortical cholinergic innervation in the human brain: Observations based on the distribution of acetylcholinesterase and choline acetyltransferase. J Comp Neurol 1988; 275: 216–240.

Mesulam M, Shaw P, Mash D, Weintraub S. Cholinergic nucleus basalis tauopathy emerges early in the aging-MCI-AD continuum. Ann Neurol 2004; 55: 815–828.

Mesulam MM. Cholinergic Innervation of Cortex by the Basal Forebrain: Cytochemistry and Cortical Connections of the Septa1 Area, Diagonal Band Nuclei, Nucleus Basalis (Substantia Innominata), and Hypothalamus in the Rhesus Monkey. J Comp Neurol 1983; 214: 170–197

Morfini G, Pigino G, Atagi Y, Busciglio J, LaDu M, Yu C, et al. Disruption of fast axonal transport is a pathogenic mechanism for intraneuronal amyloid beta. Proc Natl Acad Sci 2009; 106: 5907–5912.

Nichols E. T, Holmes P. A. Nonparametric permutation tests for functional neuroimaging: A primer with examples. Hum Brain Mapp 2002; 15: 1–25.

Nitsch RM, Blusztajn JANK, Pittast AG, Slackt BE, Growdon JH, Wurtmant RJ. Evidence for a membrane defect in Alzheimer disease brain. 1992; 89: 1671–1675.

Petersen RC, Aisen PS, Beckett LA, Donohue MC, Gamst AC, Harvey DJ, et al. Alzheimer’s Disease Neuroimaging Initiative (ADNI): Clinical characterization. Neurology 2010; 74: 201–209.

Poirier J. Apolipoprotein E in animal models of CNS injury and in alzheimer’s disease. Trends Neurosci 1994; 17: 525–530.

Poirier J, Bertrand P, Poirier J, Kogan S, Gauthier S, Poirier J, et al. Apolipoprotein E polymorphism and Alzheimer’s disease. Lancet 1993; 342: 697–699.

Poirier J, Delisle M-C, Quirion R, Aubert I, Farlow M, Lahiri D, et al. Apolipoprotein E4 allele as a predictor of cholinergic deficits and treatment outcome in Alzheimer disease. Proc Natl Acad Sci U S A 1995; 92: 12260–12264.

Roussarie J, Yao V, Plautz Z, Kasturia S, Albornoz C, Eric F, et al. Selective neuronal vulnerability in Alzheimer’ s disease: a network-based analysis. bioRxiv 2018: 1–14.

Saxena S, Caroni P. Selective Neuronal Vulnerability in Neurodegenerative Diseases: From Stressor Thresholds to Degeneration. Neuron 2011; 71: 35–48.

Saykin AJ, Shen L, Foroud TM, Potkin SG, Swaminathan S, Kim S, et al. Alzheimer’s Disease Neuroimaging Initiative biomarkers as quantitative phenotypes: Genetics core aims, progress, and plans. Alzheimer’s Dement 2010; 6: 265–273.

Schindler SE, Gray JD, Gordon BA, Xiong C, Batrla-utermann R, Quan M, et al. Cerebrospinal fluid biomarkers measured by Elecsys assays compared to amyloid imaging. Alzheimer’s Dement 2018; 14: 1460–1469.

Schmitz TW, Mur M, Aghourian M, Bedard MA, Spreng RN. Longitudinal Alzheimer’s Degeneration Reflects the Spatial Topography of Cholinergic Basal Forebrain Projections. Cell Rep 2018; 24: 38–46.

Schmitz TW, Spreng RN. Basal forebrain degeneration precedes and predicts the cortical spread of Alzheimer’s pathology. Nat Commun 2016; 7: 13249

Sepulcre J, Grothe MJ, d’Oleire Uquillas F, Ortiz-Terán L, Diez I, Yang HS, et al. Neurogenetic contributions to amyloid beta and tau spreading in the human cortex. Nat Med 2018; 24: 1910

Shaw LM, Vanderstichele H, Knapik-Czajka M, Clark CM, Aisen PS, Petersen RC, et al. Cerebrospinal fluid biomarker signature in alzheimer’s disease neuroimaging initiative subjects. Ann Neurol 2009; 65: 403–413.

Tamamaki N, Nojyo Y. Projection of the entorhinal layer II neurons in the rat as revealed by intracellular pressure-injection of neurobiotin. Hippocampus 1993; 3: 471–480.

Tan CH, Bonham LW, Fan CC, Mormino EC, Sugrue LP, Broce IJ, et al. Polygenic hazard score, amyloid deposition and Alzheimer’s neurodegeneration. Brain 2019; 142: 460–470.

Tardif CL, Collins DL, Pike GB. Regional impact of field strength on voxel-based morphometry results. Hum Brain Mapp 2010; 31: 943–957.

Veitch DP, Weiner MW, Aisen PS, Beckett LA, Cairns NJ, Green RC, et al. Understanding disease progression and improving Alzheimer’s disease clinical trials: Recent highlights from the Alzheimer’s Disease Neuroimaging Initiative. Alzheimer’s Dement 2018: 1–46.

Walker LC, Diamond MI, Duff KE, Hyman BT. Mechanisms of protein seeding in neurodegenerative diseases. JAMA Neurol 2013; 70: 304–310.

Warren JD, Rohrer JD, Schott JM, Fox NC, Hardy J, Rossor MN. Molecular nexopathies: A new paradigm of neurodegenerative disease. Trends Neurosci 2013; 36: 561–569.

Wu H, Williams J, Nathans J. Complete morphologies of basal forebrain cholinergic neurons in the mouse. Elife 2014; 2014: 1–17.

Wu JW, Hussaini SA, Bastille IM, Rodriguez GA, Mrejeru A, Rilett K, et al. Neuronal activity enhances tau propagation and tau pathology in vivo. Nat Neurosci 2016; 19: 1085–1092.

Wurtman RJ. Choline metabolism as a basis for the selective vulnerability of cholinergic neurons. Trends Neurosci 1992; 15: 117–122.

Wyman BT, Harvey DJ, Crawford K, Bernstein MA, Cole PE, Crane P, et al. Standardization of Analysis Sets for Reporting Results from ADNI MRI Data. 2014; 9: 332–337.

Zaborszky L, Hoemke L, Mohlberg H, Schleicher A, Amunts K, Zilles K. Stereotaxic probabilistic maps of the magnocellular cell groups in human basal forebrain. Neuroimage 2008; 42: 1127–1141.

