## Supplementary material for "Basal forebrain volume reliably predicts the cortical spread of Alzheimer’s degeneration"

**Short running title:** Predictive spread of Alzheimer's degeneration

**Authors:** Sara Fernández-Cabello[1,2], Martin Kronbichler[1,2,3], Koene R. A. Van Dijk[4], James A. Goodman[4], R. Nathan Spreng[5,6,7], Taylor W. Schmitz[8,9] for the Alzheimer's Disease Neuroimaging Initiative*

**Affiliations:**

[1] Department of Psychology, University of Salzburg, Salzburg, Austria

[2] Centre for Cognitive Neuroscience, University of Salzburg, Salzburg, Austria

[3] Neuroscience Institute, Christian-Doppler Medical Centre, Paracelsus Medical University, Salzburg, Austria.

[4] Clinical and Translational Imaging, Early Clinical Development, Pfizer Inc, Cambridge, MA, United States

[5] Laboratory of Brain and Cognition, Montreal Neurological Institute, Department of Neurology and Neurosurgery, McGill University, Montreal, QC, Canada

[6] Departments of Psychiatry and Psychology, McGill University, Montreal, QC, Canada

[7] Douglas Mental Health University Institute, Verdun, QC, Canada

[8] Brain and Mind Institute, Western University, London, ON, Canada

[9] Department of Physiology and Pharmacology, Western University, London, ON, Canada

* Data used in preparation of this article were obtained from the Alzheimer's Disease Neuroimaging Initiative (ADNI) database (adni.loni.usc.edu). As such, the investigators within the ADNI contributed to the design and implementation of ADNI and/or provided data but did not participate in analysis or writing of this report. A complete listing of ADNI investigators can be found at: http://adni.loni.usc.edu/wp- content/uploads/how_to_apply/ADNI_Acknowledgement_List.pdf

---

*CSF methods*

CSF samples were frozen on dry ice within 1 hour after collection and shipped overnight on dry ice to the ADNI Biomarker Core laboratory at the University of Pennsylvania Medical Center. Aliquots (0.5mL) were prepared from these and stored in barcode-labeled polypropylene vials at –80º C. The RIDs of the participants with structural MRI data were merged with the *UPENNBIOMK9_04_19_17.csv* spreadsheet to obtain within-subjects indices of baseline CSF biomarker data. The Elecsys $A\beta_{1-4}$ protocol is currently under development and the performance beyond the upper technical limit (> 1700 pg/mL) has not been formally established. These values were provided by an extrapolation of the calibration curve and are restricted to research purposes and excluded for clinical decision making.

*Rates of longitudinal degeneration by CSF groups and Clinical Diagnosis*

To investigate whether the magnitude of longitudinal degeneration in the NbM and in the EC differed between healthy individuals and individuals at different disease stages, we cross-referenced the CSF groups (abnormal CSF = pTau/Aβ >= 0.028; normal CSF = pTau/Aβ <0.028) with the ADNI baseline clinical diagnosis. This yielded 5 groups of individuals: 1) CN: normal CSF and cognitively normal; 2) MCI/AD normal CSF: normal CSF and Mild Cognitive Impairment or AD; 3) PREC: abnormal CSF and cognitively normal; 3) PROD: abnormal CSF and Mild Cognitive Impairment and 5) AD; abnormal CSF and Alzheimer's disease. See Supplementary Table 3. We then examined if the indices of longitudinal degeneration in the NbM and in the EC differed between the groups in the pooled sample (ADNI-1 and ADNI-GO/2; n=837) with a repeated-measures ANOVA covarying for Age, Gender, Education, TICV, baseline TGM, study Cohort and APOE genotype. We found a significant GROUP×ROI interaction ($F_{4,1665}$=12.94, $p$<0.001). In order to examine if the NbM and the EC dissociated at early disease stages, we focused the post-hoc comparison in the CN and the PREC groups. Independent sample t-tests revealed that the PREC group had larger magnitudes of degeneration in the NbM ($t_{118}$=2.27, $p$=0.025). However, and inconsistent with our previous report (Schmitz and Spreng, 2016), the magnitude of degeneration in the EC was also more pronounced in the PREC group than in the CN ($t_{108}$=2.70, $p$=0.007). In addition, we found a main effect of Group ($F_{4,1665}$=13.30, $p$<0.001) and ROI ($F_{4,1665}$=12.70, $p$<0.001). See Supplementary Fig. 2.

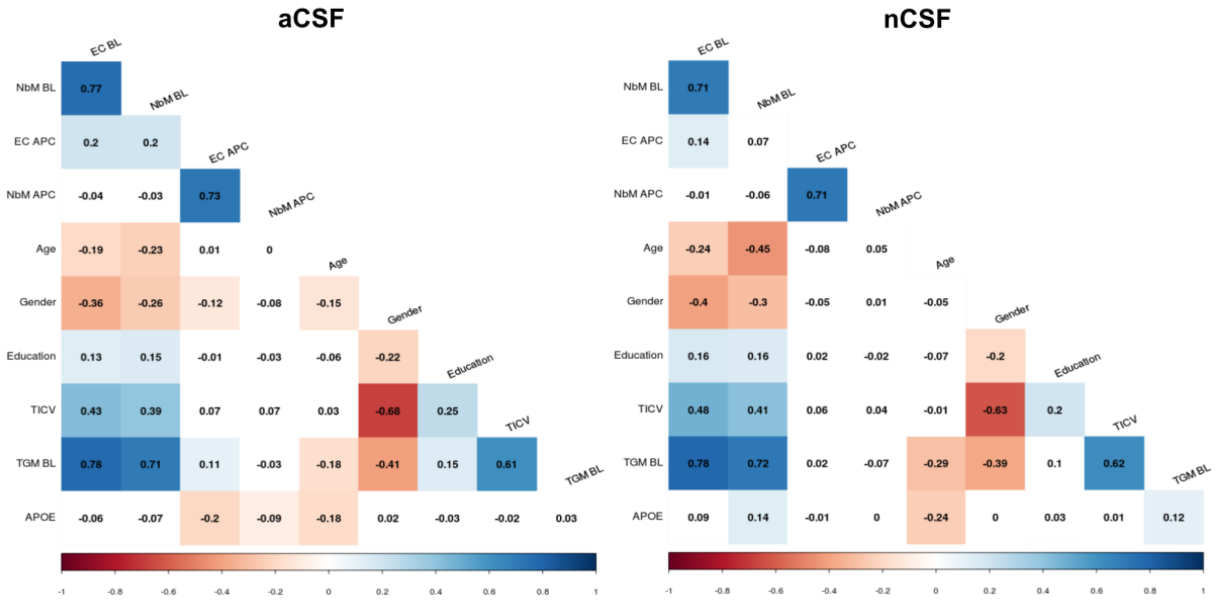


**Supplementary Figure 1. Association between the variables included in the robust regression models.** Pearson's correlation coefficients between the variables included in the robust regression models of the pooled (ADNI-1 and ADNI-GO/2) CSF groups. Note that the covariates (Age, Gender, Education, TICV and baseline TGM) are not statistically independent from the predictor variables (baseline volumes). Annual percent change (APC) volumes are the predicted variables in all the models. Colored squares denote statistically significant correlations at $p<0.05$. aCSF=abnormal CSF group; nCSF=normal CSF group; NbM=nucleus basalis of Meynert; EC=entorhinal cortex; BL=Baseline; APC=annual percent change; TICV=total intracranial volume; TGM=total gray matter.

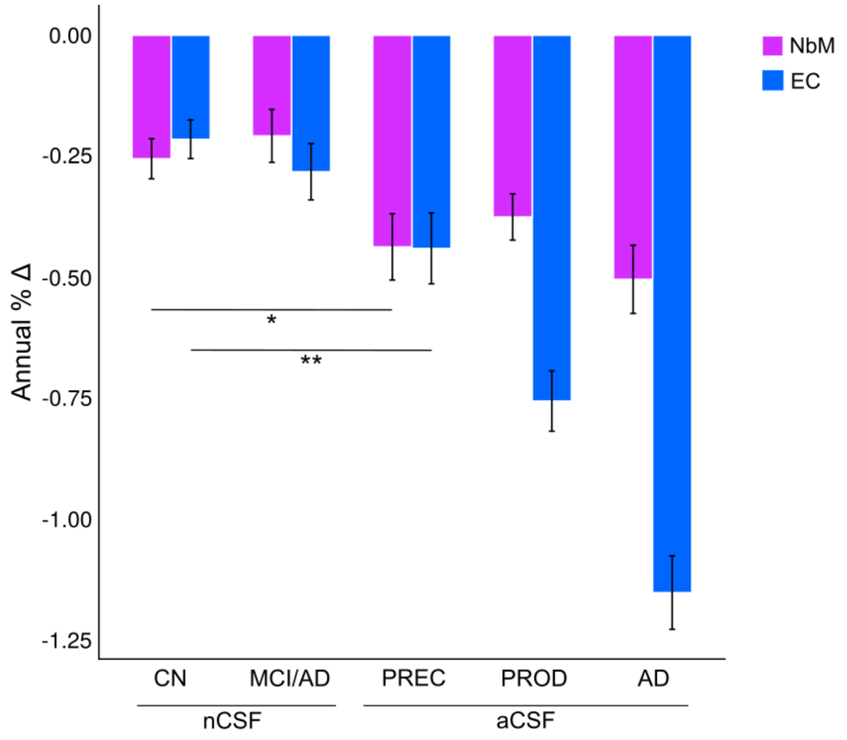


**Supplementary Figure 2. GM degeneration according to CSF Group and Clinical Diagnosis.** The magnitudes of longitudinal degeneration in the NbM and the EC differed between groups with different CSF status (normal vs. abnormal) and baseline clinical diagnosis. In the early disease stages (preclinical), NbM and EC degeneration was more pronounced than in the healthy control group (cognitively normal). CN=cognitively normal; MCI/AD=mild cognitively impaired or Alzheimer's disease; PREC=preclinical; PROD=prodromal; AD=Alzheimer's disease; aCSF=abnormal CSF group; nCSF=normal CSF group; NbM=nucleus basalis of Meynert; EC = entorhinal cortex.

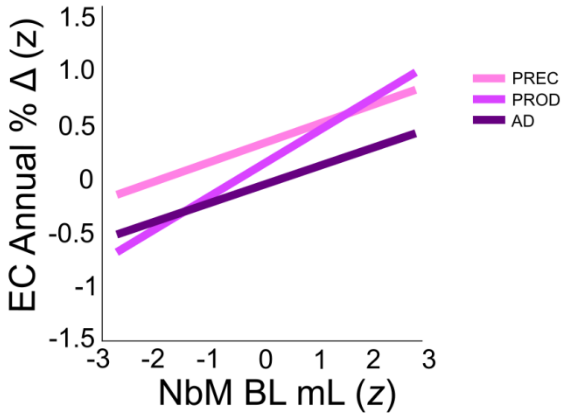


**Supplementary Figure 3. Moderation of Clinical Diagnosis.** The predictive relationship between baseline NbM and longitudinal degeneration in the EC was not moderated by the ADNI baseline Clinical Diagnosis (see main text). PREC=preclinical; PROD=prodromal; AD=Alzheimer's disease. The x axis reflects baseline NbM volumes and the y axis reflects annual percent change in the EC. Baseline volumes and annual percent change units were z-scored (z). NbM=nucleus basalis of Meynert; EC=entorhinal cortex; mL=milliliters; BL=baseline volume.

**Supplementary Table 1**

*Participant Demographics by CSF Group and Baseline Clinical Diagnosis*

| | CSF group | | | | |
|---|---|---|---|---|---|
| | **nCSF** | | **aCSF** | | |
| | CN | MCI/AD | PREC | PROD | AD |
| N | 226 | 121 | 67 | 219 | 204 |
| Age | 71.74 (6.21) | 72.63 (8.15) | 74.97 (5.27) | 72.88 (7.21) | 72.46 (7.72) |
| Female/Male | 118/108 | 42/79 | 38/29 | 85/134 | 89/115 |
| Education | 16.33 (2.64) | 16.5 (2.77) | 15.81 (2.82) | 15.87 (2.92) | 15.77 (2.62) |
| ε4+/ε4- | 44/182 | 32/89 | 35/32 | 152/67 | 159/45 |
| $A\beta_{1\text{-}42}$ | 1574.98 (579.71) | 1089.76 (428.65) | 643.99 (229.93) | 630.99 (179.54) | 599.14 (185.96) |
| pTau | 19.47 (6.50) | 17.16 (6.31) | 29.41 (10.1) | 35.14 (14.67) | 37.28 (14.12) |

Information of the included participants from the pooled (ADNI-1 and ADNI-GO/2) sample by CSF group and baseline Clinical Diagnosis. Values represent means (standard deviations) unless otherwise specified. aCSF=abnormal CSF group; nCSF=normal CSF group; CN=Cognitively Normal; PREC=preclinical; PROD=prodromal; AD=Alzheimer's Disease. Aβ=CSF concentration of the amyloid beta 1-42 peptide; pTau=CSF concentration of hyper phosphorylated tau (at threonine 181). APOE status: ε4+ = ε4 carrier, ε4- = ε4 non-carrier. Age and Education are in years. Aβ1-42 and pTau concentrations are in picograms per milliliter (pg/mL). Values in parentheses are standard deviations.

### Supplementary Table 2

*Output of the Robust Regression Models in the Pooled (ADNI-1 and ADNI-GO/2) Sample*

| | CSF Group | | | | | |
|---|---|---|---|---|---|---|
| | **aCSF** (n=490) NbM→EC | | | **nCSF** (n=347) NbM→EC | | |
| | Estimate | SE | t | Estimate | SE | t |
| Intercept | 0.32 | 0.07 | 4.68*** | 0.09 | 0.05 | 1.87 |
| Age | 0.07 | 0.04 | 1.79 | -0.09 | 0.05 | -1.73 |
| Female/Male | -0.05 | 0.05 | -1.06 | -0.02 | 0.06 | -0.41 |
| Education | -0.01 | 0.04 | -0.23 | 0.02 | 0.05 | 0.53 |
| TICV | -0.06 | 0.06 | -1.00 | 0.06 | 0.07 | 0.88 |
| TGM Baseline | -0.08 | 0.06 | -1.27 | -0.14 | 0.08 | -1.77 |
| Cohort | -0.13 | 0.04 | -3.33*** | -0.10 | 0.05 | -1.92 |
| ε4+/ε4- | -0.29 | 0.08 | -3.56*** | -0.04 | 0.11 | -0.37 |
| NbM Baseline | 0.30 | 0.06 | 5.32*** | 0.11 | 0.08 | 1.45 |
| | R-Squared: 0.103; F-statistic vs. constant model: 2.73, p-value = 0.0105 | | | R-squared: 0.0366; F-statistic vs. constant model: 1.61, p-value = 0.122 | | |

| | EC→NbM | | | EC→NbM | | |
|---|---|---|---|---|---|---|
| | Estimate | SE | T | Estimate | SE | t |
| Intercept | 0.16 | 0.07 | 2.15** | 0.08 | 0.05 | 1.52 |
| Age | -0.03 | 0.04 | -0.74 | 0.01 | 0.05 | 0.17 |
| Female/Male | -0.08 | 0.06 | -1.39 | 0.05 | 0.06 | 0.76 |
| Education | -0.01 | 0.04 | -0.16 | 0.02 | 0.05 | 0.33 |
| TICV | 0.04 | 0.06 | 0.61 | 0.14 | 0.07 | 1.96 |
| TGM Baseline | -0.10 | 0.07 | -1.42 | -0.22 | 0.09 | -2.46** |
| Cohort | -0.15 | 0.04 | -3.74*** | -0.05 | 0.05 | -0.94 |
| ε4+/ε4- | -0.17 | 0.09 | -1.90 | 0.00 | 0.12 | 0.01 |
| EC Baseline | 0.02 | 0.07 | 0.26 | 0.12 | 0.08 | 1.53 |
| | R-Squared: 0.052; F-statistic vs. constant model: 3.36, p-value = 0.000925 | | | R-squared: 0.035; F-statistic vs. constant model: 1.53, p-value = 0.145 | | |

Estimated coefficients of the robust linear regression models in the pooled (ADNI-1 and ADNI-GO/2) sample. For each CSF group, two regression models were fitted (NbM→EC and EC→NbM). Values represent the estimated coefficients (Estimate), their standard errors (SE) and the associated t statistic (t) of the predictor variables. aCSF=abnormal CSF group; nCSF=normal CSF group; NbM=nucleus basalis of Meynert, EC=entorhinal cortex; TIVC=total intracranial volume; TGM=total gray matter; APOE status: ε4+ = ε4 carrier, ε4- = ε4 non-carrier. ***$p$<0.001, **$p$<0.005.

### Supplementary Table 3

*Robust Regression Models without Covariates*

| | | CSF Group | | | | | | | | | | | |
|---|---|---|---|---|---|---|---|---|---|---|---|---|---|
| | | **aCSF** | | | | | | **nCSF** | | | | | |
| **Dataset** | | NbM→EC | | | EC→NbM | | | NbM→EC | | | EC→NbM | | |
| | | Estimate | SE | t | Estimate | SE | t | Estimate | SE | t | Estimate | SE | t |
| **ADNI-1** | Intercept | 0.13 | 0.05 | 2.35** | 0.06 | 0.06 | 0.95 | 0.08 | 0.08 | 0.93 | 0.13 | 0.08 | 1.54 |
| | Baseline Volume | 0.19 | 0.05 | 3.43*** | 0.00 | 0.06 | 0.06 | -0.00 | 0.08 | -0.08 | 0.08 | 0.08 | 0.99 |
| | | R-squared=0.09; p<0.001 | | | R-squared=0.00; p=0.34 | | | R-squared=0.008; p=0.35 | | | R-squared=0.03; p=0.06 | | |
| | | NbM→EC | | | EC→NbM | | | NbM→EC | | | EC→NbM | | |
| | | Estimate | SE | t | Estimate | SE | t | Estimate | SE | t | Estimate | SE | t |
| **ADNI-GO/2** | Intercept | 0.9 | 0.04 | 2.06** | 0.02 | 0.04 | 0.53 | 0.08 | 0.05 | 1.76 | 0.08 | 0.5 | 1.47 |
| | Baseline Volume | 0.24 | 0.04 | 5.23*** | 0.00 | 0.04 | 0.03 | 0.09 | 0.05 | $1.90^{+}$ | -0.03 | 0.05 | -0.63 |
| | | R-squared=0.09; p<0.001 | | | R-squared=0.00; p=0.59 | | | R-squared=0.03; p=0.01; $^{+}p = 0.058$ | | | R-squared=0.01; p=0.11 | | |
| | | NbM→EC | | | EC→NbM | | | NbM→EC | | | EC→NbM | | |
| | | Estimate | SE | t | Estimate | SE | t | Estimate | SE | t | Estimate | SE | t |
| **Pooled** | Intercept | 0.13 | 0.03 | 3.71*** | 0.06 | 0.03 | 1.63 | 0.10 | 0.04 | 2.46** | 0.08 | 0.04 | 2.01** |
| | Baseline Volume | 0.18 | 0.03 | 5.15*** | -0.04 | 0.03 | -1.28 | 0.03 | 0.04 | 0.94 | -0.01 | 0.044 | -0.17 |
| | | R-squared=0.07; p<0.001 | | | R-squared=0.00; p=0.03 | | | R-squared=0.02; p=0.01 | | | R-squared=0.01; p=0.04 | | |

To ensure that our main findings were not distorted by collinearities in the model predictors, we re-estimated each model without the covariates for age, sex, education, TICV and baseline TGM. Baseline NbM volumes predicted longitudinal degeneration in the EC in all datasets. Values represent the estimated coefficients (Estimate), their standard errors (SE) and the associated t statistic (t) of the predictor variables. For each CSF group, two regression models were fitted (NbM→EC and EC→NbM). For the NbM→EC models, “Baseline Volume” refers to NbM. For the EC→NbM, “Baseline Volume” refers to EC. aCSF=abnormal CSF group; nCSF=normal CSF group; NbM=nucleus basalis of Meynert; EC=entorhinal cortex. ***$p<0.001$, **$p<0.005$.

**Supplementary Table 4**

*Study Participants*

| RID | Cohort | pTau | A$\beta_{1-42}$ | pTau/A$\beta_{1-42}$ | CSF Group |
|---|---|---|---|---|---|
| *S_0003 | ADNI1 | 22.83 | 741.5 | 0.031 | aCSF |
| *S_0005 | ADNI1 | 33.43 | 547.3 | 0.061 | aCSF |
| *S_0010 | ADNI1 | 31.26 | 357.4 | 0.087 | aCSF |
| *S_0014 | ADNI1 | 16.68 | 1582 | 0.011 | nCSF |
| *S_0016 | ADNI1 | 15.88 | 550.6 | 0.029 | aCSF |
| *S_0022 | ADNI1 | 21.85 | 2438 | 0.009 | nCSF |
| *S_0023 | ADNI1 | 16.74 | 1647 | 0.010 | nCSF |
| *S_0029 | ADNI1 | 27.24 | 442.9 | 0.062 | aCSF |
| *S_0031 | ADNI1 | 22.55 | 1774 | 0.013 | nCSF |
| *S_0033 | ADNI1 | 34.93 | 673 | 0.052 | aCSF |
| *S_0040 | ADNI1 | 12.5 | 1526 | 0.008 | nCSF |
| *S_0041 | ADNI1 | 18.07 | 414.6 | 0.044 | aCSF |
| *S_0051 | ADNI1 | 28.4 | 352.5 | 0.081 | aCSF |
| *S_0057 | ADNI1 | 30.25 | 597.1 | 0.051 | aCSF |
| *S_0061 | ADNI1 | 22.08 | 1288 | 0.017 | nCSF |
| *S_0066 | ADNI1 | 15.97 | 1477 | 0.011 | nCSF |
| *S_0076 | ADNI1 | 44.86 | 503.5 | 0.089 | aCSF |
| *S_0077 | ADNI1 | 65.4 | 792.7 | 0.083 | aCSF |
| *S_0083 | ADNI1 | 45.92 | 314.7 | 0.146 | aCSF |
| *S_0084 | ADNI1 | 29.85 | 479.6 | 0.062 | aCSF |
| *S_0086 | ADNI1 | 15.79 | 950.1 | 0.017 | nCSF |
| *S_0089 | ADNI1 | 14.88 | 1414 | 0.011 | nCSF |
| *S_0091 | ADNI1 | 30.16 | 502.6 | 0.060 | aCSF |
| *S_0093 | ADNI1 | 20.76 | 562.4 | 0.037 | aCSF |
| *S_0094 | ADNI1 | 36.96 | 462.4 | 0.080 | aCSF |
| *S_0096 | ADNI1 | 12.17 | 915.3 | 0.013 | nCSF |
| *S_0097 | ADNI1 | 18.06 | 776.6 | 0.023 | nCSF |
| *S_0101 | ADNI1 | 14.62 | 453.2 | 0.032 | aCSF |
| *S_0102 | ADNI1 | 27.85 | 539.3 | 0.052 | aCSF |
| *S_0106 | ADNI1 | 18.52 | 431.4 | 0.043 | aCSF |
| *S_0107 | ADNI1 | 16.31 | 1652 | 0.010 | nCSF |
| *S_0112 | ADNI1 | 24.43 | 498.6 | 0.049 | aCSF |
| *S_0118 | ADNI1 | 16.23 | 1467 | 0.011 | nCSF |
| *S_0120 | ADNI1 | 14.38 | 1226 | 0.012 | nCSF |
| *S_0123 | ADNI1 | 26.3 | 1566 | 0.017 | nCSF |
| *S_0126 | ADNI1 | 24.94 | 536.8 | 0.046 | aCSF |

| | | | | | |
|---|---|---|---|---|---|
| *S_0127 | ADNI1 | 30.66 | 2573 | 0.012 | nCSF |
| *S_0129 | ADNI1 | 26.45 | 682.7 | 0.039 | aCSF |
| *S_0130 | ADNI1 | 26.53 | 509.7 | 0.052 | aCSF |
| *S_0139 | ADNI1 | 45.14 | 651.9 | 0.069 | aCSF |
| *S_0147 | ADNI1 | 23.33 | 371.4 | 0.063 | aCSF |
| *S_0149 | ADNI1 | 19.07 | 652.1 | 0.029 | aCSF |
| *S_0150 | ADNI1 | 47.6 | 817 | 0.058 | aCSF |
| *S_0158 | ADNI1 | 21.69 | 2056 | 0.011 | nCSF |
| *S_0159 | ADNI1 | 9.74 | 622.7 | 0.016 | nCSF |
| *S_0172 | ADNI1 | 21.19 | 2070 | 0.010 | nCSF |
| *S_0173 | ADNI1 | 29.3 | 991.3 | 0.030 | aCSF |
| *S_0176 | ADNI1 | 40.9 | 670.7 | 0.061 | aCSF |
| *S_0177 | ADNI1 | 21.41 | 1367 | 0.016 | nCSF |
| *S_0186 | ADNI1 | 16.77 | 942.8 | 0.018 | nCSF |
| *S_0196 | ADNI1 | 16.32 | 540.6 | 0.030 | aCSF |
| *S_0204 | ADNI1 | 20.29 | 282.9 | 0.072 | aCSF |
| *S_0210 | ADNI1 | 18.54 | 1385 | 0.013 | nCSF |
| *S_0214 | ADNI1 | 48.14 | 683.1 | 0.070 | aCSF |
| *S_0221 | ADNI1 | 44.39 | 379.1 | 0.117 | aCSF |
| *S_0222 | ADNI1 | 31.9 | 620.3 | 0.051 | aCSF |
| *S_0223 | ADNI1 | 41.03 | 751.5 | 0.055 | aCSF |
| *S_0231 | ADNI1 | 31.15 | 426.1 | 0.073 | aCSF |
| *S_0232 | ADNI1 | 19.67 | 1028 | 0.019 | nCSF |
| *S_0240 | ADNI1 | 22.91 | 1338 | 0.017 | nCSF |
| *S_0256 | ADNI1 | 27.12 | 288.4 | 0.094 | aCSF |
| *S_0257 | ADNI1 | 21.4 | 427.2 | 0.050 | aCSF |
| *S_0259 | ADNI1 | 32.32 | 803.6 | 0.040 | aCSF |
| *S_0260 | ADNI1 | 15.47 | 1520 | 0.010 | nCSF |
| *S_0269 | ADNI1 | 32.75 | 576.8 | 0.057 | aCSF |
| *S_0273 | ADNI1 | 16.11 | 1639 | 0.010 | nCSF |
| *S_0285 | ADNI1 | 37.99 | 844.4 | 0.045 | aCSF |
| *S_0286 | ADNI1 | 40.28 | 642.9 | 0.063 | aCSF |
| *S_0291 | ADNI1 | 38.55 | 802.6 | 0.048 | aCSF |
| *S_0292 | ADNI1 | 13.1 | 941.7 | 0.014 | nCSF |
| *S_0293 | ADNI1 | 45.23 | 702.1 | 0.064 | aCSF |
| *S_0294 | ADNI1 | 24.75 | 2387 | 0.010 | nCSF |
| *S_0295 | ADNI1 | 34.73 | 888.1 | 0.039 | aCSF |
| *S_0299 | ADNI1 | 12.8 | 997.1 | 0.013 | nCSF |
| *S_0300 | ADNI1 | 32.87 | 475.2 | 0.069 | aCSF |
| *S_0307 | ADNI1 | 11.99 | 942.5 | 0.013 | nCSF |
| *S_0314 | ADNI1 | 32.43 | 379.7 | 0.085 | aCSF |

| | | | | | |
|---|---|---|---|---|---|
| *S_0316 | ADNI1 | 43.44 | 300.9 | 0.144 | aCSF |
| *S_0326 | ADNI1 | 24.04 | 1876 | 0.013 | nCSF |
| *S_0327 | ADNI1 | 16.71 | 1405 | 0.012 | nCSF |
| *S_0331 | ADNI1 | 39.86 | 588 | 0.068 | aCSF |
| *S_0336 | ADNI1 | 45.17 | 854.2 | 0.053 | aCSF |
| *S_0341 | ADNI1 | 19.33 | 313.1 | 0.062 | aCSF |
| *S_0344 | ADNI1 | 29.26 | 556.7 | 0.053 | aCSF |
| *S_0352 | ADNI1 | 15.45 | 1239 | 0.012 | nCSF |
| *S_0361 | ADNI1 | 38.22 | 585.5 | 0.065 | aCSF |
| *S_0362 | ADNI1 | 78.46 | 414.1 | 0.189 | aCSF |
| *S_0366 | ADNI1 | 30.61 | 508.9 | 0.060 | aCSF |
| *S_0372 | ADNI1 | 28.66 | 967 | 0.030 | aCSF |
| *S_0374 | ADNI1 | 24.03 | 495.2 | 0.049 | aCSF |
| *S_0376 | ADNI1 | 13.21 | 1148 | 0.012 | nCSF |
| *S_0378 | ADNI1 | 56.81 | 482.9 | 0.118 | aCSF |
| *S_0386 | ADNI1 | 16.77 | 1168 | 0.014 | nCSF |
| *S_0388 | ADNI1 | 31.59 | 332.6 | 0.095 | aCSF |
| *S_0390 | ADNI1 | 59.84 | 490 | 0.122 | aCSF |
| *S_0394 | ADNI1 | 33.87 | 643.4 | 0.053 | aCSF |
| *S_0400 | ADNI1 | 28.51 | 265.6 | 0.107 | aCSF |
| *S_0403 | ADNI1 | 36.23 | 333.5 | 0.109 | aCSF |
| *S_0404 | ADNI1 | 42.87 | 2401 | 0.018 | nCSF |
| *S_0413 | ADNI1 | 10.57 | 1006 | 0.011 | nCSF |
| *S_0422 | ADNI1 | 21.84 | 1907 | 0.011 | nCSF |
| *S_0424 | ADNI1 | 17.94 | 399.1 | 0.045 | aCSF |
| *S_0426 | ADNI1 | 14.3 | 374.5 | 0.038 | aCSF |
| *S_0429 | ADNI1 | 15.55 | 1014 | 0.015 | nCSF |
| *S_0431 | ADNI1 | 38.61 | 657.2 | 0.059 | aCSF |
| *S_0433 | ADNI1 | 24.9 | 831.3 | 0.030 | aCSF |
| *S_0441 | ADNI1 | 21.45 | 1614 | 0.013 | nCSF |
| *S_0443 | ADNI1 | 27.73 | 1850 | 0.015 | nCSF |
| *S_0448 | ADNI1 | 20.87 | 1645 | 0.013 | nCSF |
| *S_0454 | ADNI1 | 31.71 | 2108 | 0.015 | nCSF |
| *S_0457 | ADNI1 | 34.79 | 471.5 | 0.074 | aCSF |
| *S_0459 | ADNI1 | 27.12 | 2490 | 0.011 | nCSF |
| *S_0464 | ADNI1 | 14.12 | 1203 | 0.012 | nCSF |
| *S_0467 | ADNI1 | 23.98 | 500.5 | 0.048 | aCSF |
| *S_0470 | ADNI1 | 44.28 | 725.2 | 0.061 | aCSF |
| *S_0472 | ADNI1 | 20.22 | 1550 | 0.013 | nCSF |
| *S_0474 | ADNI1 | 41.6 | 441.2 | 0.094 | aCSF |
| *S_0479 | ADNI1 | 22.8 | 649.7 | 0.035 | aCSF |

| | | | | | |
|---|---|---|---|---|---|
| *S_0481 | ADNI1 | 35.95 | 594 | 0.061 | aCSF |
| *S_0488 | ADNI1 | 17.22 | 873.8 | 0.020 | nCSF |
| *S_0498 | ADNI1 | 19.71 | 2001 | 0.010 | nCSF |
| *S_0507 | ADNI1 | 31.48 | 449.8 | 0.070 | aCSF |
| *S_0511 | ADNI1 | 35.77 | 627.5 | 0.057 | aCSF |
| *S_0514 | ADNI1 | 10.41 | 826.8 | 0.013 | nCSF |
| *S_0516 | ADNI1 | 29.6 | 1860 | 0.016 | nCSF |
| *S_0518 | ADNI1 | 29.13 | 540.2 | 0.054 | aCSF |
| *S_0519 | ADNI1 | 15.07 | 1263 | 0.012 | nCSF |
| *S_0520 | ADNI1 | 20.72 | 600.2 | 0.035 | aCSF |
| *S_0533 | ADNI1 | 12.91 | 637.8 | 0.020 | nCSF |
| *S_0534 | ADNI1 | 12.95 | 806.8 | 0.016 | nCSF |
| *S_0535 | ADNI1 | 32.44 | 623 | 0.052 | aCSF |
| *S_0543 | ADNI1 | 44.35 | 663.7 | 0.067 | aCSF |
| *S_0544 | ADNI1 | 12.46 | 461.6 | 0.027 | nCSF |
| *S_0547 | ADNI1 | 32.4 | 2568 | 0.013 | nCSF |
| *S_0552 | ADNI1 | 39.15 | 648.4 | 0.060 | aCSF |
| *S_0555 | ADNI1 | 20.13 | 560.6 | 0.036 | aCSF |
| *S_0559 | ADNI1 | 15.82 | 1245 | 0.013 | nCSF |
| *S_0565 | ADNI1 | 27.29 | 212.3 | 0.129 | aCSF |
| *S_0566 | ADNI1 | 29.77 | 682.5 | 0.044 | aCSF |
| *S_0567 | ADNI1 | 20.69 | 368.6 | 0.056 | aCSF |
| *S_0577 | ADNI1 | 55.22 | 702.2 | 0.079 | aCSF |
| *S_0578 | ADNI1 | 27.56 | 585.6 | 0.047 | aCSF |
| *S_0579 | ADNI1 | 19.43 | 1429 | 0.014 | nCSF |
| *S_0588 | ADNI1 | 40.69 | 529.2 | 0.077 | aCSF |
| *S_0602 | ADNI1 | 16.98 | 1251 | 0.014 | nCSF |
| *S_0604 | ADNI1 | 30.47 | 1079 | 0.028 | aCSF |
| *S_0605 | ADNI1 | 23.7 | 1117 | 0.021 | nCSF |
| *S_0607 | ADNI1 | 39.89 | 507.3 | 0.079 | aCSF |
| *S_0610 | ADNI1 | 32.15 | 3187 | 0.010 | nCSF |
| *S_0618 | ADNI1 | 18.66 | 1323 | 0.014 | nCSF |
| *S_0619 | ADNI1 | 18.56 | 393.8 | 0.047 | aCSF |
| *S_0621 | ADNI1 | 42.85 | 467 | 0.092 | aCSF |
| *S_0625 | ADNI1 | 20.33 | 645.9 | 0.031 | aCSF |
| *S_0626 | ADNI1 | 36.67 | 644.9 | 0.057 | aCSF |
| *S_0634 | ADNI1 | 34.68 | 1770 | 0.020 | nCSF |
| *S_0637 | ADNI1 | 26.17 | 2594 | 0.010 | nCSF |
| *S_0638 | ADNI1 | 23.74 | 269.3 | 0.088 | aCSF |
| *S_0644 | ADNI1 | 16.04 | 338.4 | 0.047 | aCSF |
| *S_0648 | ADNI1 | 25.18 | 1479 | 0.017 | nCSF |

| | | | | | |
|---|---|---|---|---|---|
| *S_0649 | ADNI1 | 42.03 | 669.3 | 0.063 | aCSF |
| *S_0657 | ADNI1 | 15.06 | 1697 | 0.009 | nCSF |
| *S_0658 | ADNI1 | 35.69 | 488.5 | 0.073 | aCSF |
| *S_0671 | ADNI1 | 35.23 | 578.9 | 0.061 | aCSF |
| *S_0672 | ADNI1 | 19.13 | 1631 | 0.012 | nCSF |
| *S_0673 | ADNI1 | 62.83 | 455.1 | 0.138 | aCSF |
| *S_0677 | ADNI1 | 15.07 | 988.7 | 0.015 | nCSF |
| *S_0680 | ADNI1 | 19.69 | 1868 | 0.011 | nCSF |
| *S_0685 | ADNI1 | 22.82 | 2296 | 0.010 | nCSF |
| *S_0686 | ADNI1 | 13.56 | 657.6 | 0.021 | nCSF |
| *S_0690 | ADNI1 | 30.8 | 756.9 | 0.041 | aCSF |
| *S_0692 | ADNI1 | 8 | 390.9 | 0.020 | nCSF |
| *S_0717 | ADNI1 | 40.02 | 494.3 | 0.081 | aCSF |
| *S_0722 | ADNI1 | 24.96 | 1954 | 0.013 | nCSF |
| *S_0723 | ADNI1 | 18.02 | 692.3 | 0.026 | nCSF |
| *S_0724 | ADNI1 | 42.12 | 812.7 | 0.052 | aCSF |
| *S_0725 | ADNI1 | 21.53 | 566.7 | 0.038 | aCSF |
| *S_0729 | ADNI1 | 31.69 | 957.8 | 0.033 | aCSF |
| *S_0731 | ADNI1 | 27.56 | 744.2 | 0.037 | aCSF |
| *S_0733 | ADNI1 | 45.2 | 711.9 | 0.063 | aCSF |
| *S_0739 | ADNI1 | 39.06 | 1066 | 0.037 | aCSF |
| *S_0746 | ADNI1 | 22.1 | 1837 | 0.012 | nCSF |
| *S_0748 | ADNI1 | 16.76 | 520.8 | 0.032 | aCSF |
| *S_0750 | ADNI1 | 53.41 | 533.1 | 0.100 | aCSF |
| *S_0753 | ADNI1 | 32.08 | 547.6 | 0.059 | aCSF |
| *S_0754 | ADNI1 | 34.11 | 392.1 | 0.087 | aCSF |
| *S_0768 | ADNI1 | 18.85 | 404.3 | 0.047 | aCSF |
| *S_0778 | ADNI1 | 20.65 | 668.2 | 0.031 | aCSF |
| *S_0779 | ADNI1 | 28.19 | 742 | 0.038 | aCSF |
| *S_0783 | ADNI1 | 22.19 | 1038 | 0.021 | nCSF |
| *S_0784 | ADNI1 | 13.46 | 476.1 | 0.028 | aCSF |
| *S_0796 | ADNI1 | 13.11 | 1071 | 0.012 | nCSF |
| *S_0800 | ADNI1 | 22.75 | 565.2 | 0.040 | aCSF |
| *S_0803 | ADNI1 | 70.52 | 1147 | 0.061 | aCSF |
| *S_0814 | ADNI1 | 49.55 | 749.9 | 0.066 | aCSF |
| *S_0818 | ADNI1 | 18.7 | 1843 | 0.010 | nCSF |
| *S_0834 | ADNI1 | 33.9 | 524.4 | 0.065 | aCSF |
| *S_0835 | ADNI1 | 28.62 | 562.6 | 0.051 | aCSF |
| *S_0839 | ADNI1 | 44.42 | 730.1 | 0.061 | aCSF |
| *S_0843 | ADNI1 | 11.86 | 1298 | 0.009 | nCSF |
| *S_0850 | ADNI1 | 31.73 | 618.2 | 0.051 | aCSF |

| | | | | | |
|---|---|---|---|---|---|
| *S_0861 | ADNI1 | 30.85 | 371.4 | 0.083 | aCSF |
| *S_0866 | ADNI1 | 16.13 | 1271 | 0.013 | nCSF |
| *S_0878 | ADNI1 | 37.97 | 526.4 | 0.072 | aCSF |
| *S_0886 | ADNI1 | 17.65 | 1737 | 0.010 | nCSF |
| *S_0891 | ADNI1 | 22.49 | 680.8 | 0.033 | aCSF |
| *S_0896 | ADNI1 | 17.58 | 1208 | 0.015 | nCSF |
| *S_0904 | ADNI1 | 50.97 | 573.6 | 0.089 | aCSF |
| *S_0906 | ADNI1 | 32.63 | 908.9 | 0.036 | aCSF |
| *S_0908 | ADNI1 | 14.64 | 1314 | 0.011 | nCSF |
| *S_0912 | ADNI1 | 32.75 | 2079 | 0.016 | nCSF |
| *S_0921 | ADNI1 | 48.22 | 608.2 | 0.079 | aCSF |
| *S_0923 | ADNI1 | 20.83 | 1396 | 0.015 | nCSF |
| *S_0925 | ADNI1 | 12.15 | 602.8 | 0.020 | nCSF |
| *S_0926 | ADNI1 | 14.22 | 1351 | 0.011 | nCSF |
| *S_0932 | ADNI1 | 22.57 | 579.2 | 0.039 | aCSF |
| *S_0941 | ADNI1 | 28.06 | 755.1 | 0.037 | aCSF |
| *S_0950 | ADNI1 | 25.21 | 494 | 0.051 | aCSF |
| *S_0952 | ADNI1 | 19.45 | 385.4 | 0.050 | aCSF |
| *S_0961 | ADNI1 | 55.17 | 313.5 | 0.176 | aCSF |
| *S_0972 | ADNI1 | 46.32 | 696 | 0.067 | aCSF |
| *S_0973 | ADNI1 | 18.33 | 1604 | 0.011 | nCSF |
| *S_0978 | ADNI1 | 37.9 | 657.6 | 0.058 | aCSF |
| *S_0981 | ADNI1 | 34.13 | 1606 | 0.021 | nCSF |
| *S_0984 | ADNI1 | 36.02 | 200 | 0.180 | aCSF |
| *S_0994 | ADNI1 | 51.42 | 699.3 | 0.074 | aCSF |
| *S_0997 | ADNI1 | 40.44 | 944.8 | 0.043 | aCSF |
| *S_1002 | ADNI1 | 12.57 | 1449 | 0.009 | nCSF |
| *S_1009 | ADNI1 | 21.39 | 561.3 | 0.038 | aCSF |
| *S_1010 | ADNI1 | 27.78 | 594.9 | 0.047 | aCSF |
| *S_1016 | ADNI1 | 24.71 | 1044 | 0.024 | nCSF |
| *S_1034 | ADNI1 | 15.79 | 666.2 | 0.024 | nCSF |
| *S_1035 | ADNI1 | 34.21 | 935.3 | 0.037 | aCSF |
| *S_1041 | ADNI1 | 28.22 | 502.4 | 0.056 | aCSF |
| *S_1045 | ADNI1 | 19.72 | 2037 | 0.010 | nCSF |
| *S_1046 | ADNI1 | 17.92 | 732.9 | 0.024 | nCSF |
| *S_1054 | ADNI1 | 39.09 | 511.9 | 0.076 | aCSF |
| *S_1063 | ADNI1 | 28.29 | 685.5 | 0.041 | aCSF |
| *S_1073 | ADNI1 | 69.63 | 657.9 | 0.106 | aCSF |
| *S_1081 | ADNI1 | 37.49 | 596 | 0.063 | aCSF |
| *S_1082 | ADNI1 | 32.96 | 517.6 | 0.064 | aCSF |
| *S_1097 | ADNI1 | 50.89 | 775.6 | 0.066 | aCSF |

| | | | | | |
|---|---|---|---|---|---|
| *S_1098 | ADNI1 | 48.27 | 561.7 | 0.086 | aCSF |
| *S_1109 | ADNI1 | 32.4 | 449.2 | 0.072 | aCSF |
| *S_1126 | ADNI1 | 22.82 | 566.3 | 0.040 | aCSF |
| *S_1130 | ADNI1 | 56.14 | 664.3 | 0.085 | aCSF |
| *S_1140 | ADNI1 | 19.71 | 1735 | 0.011 | nCSF |
| *S_1144 | ADNI1 | 35.2 | 564.2 | 0.062 | aCSF |
| *S_1168 | ADNI1 | 19.88 | 915 | 0.022 | nCSF |
| *S_1169 | ADNI1 | 34.78 | 3357 | 0.010 | nCSF |
| *S_1170 | ADNI1 | 48.82 | 730.9 | 0.067 | aCSF |
| *S_1171 | ADNI1 | 15.13 | 511.1 | 0.030 | aCSF |
| *S_1183 | ADNI1 | 30.47 | 1583 | 0.019 | nCSF |
| *S_1187 | ADNI1 | 26.44 | 1714 | 0.015 | nCSF |
| *S_1190 | ADNI1 | 33.74 | 1880 | 0.018 | nCSF |
| *S_1206 | ADNI1 | 20.77 | 1979 | 0.010 | nCSF |
| *S_1213 | ADNI1 | 37.44 | 574 | 0.065 | aCSF |
| *S_1217 | ADNI1 | 23.8 | 912 | 0.026 | nCSF |
| *S_1224 | ADNI1 | 26.56 | 496.4 | 0.054 | aCSF |
| *S_1227 | ADNI1 | 18.66 | 1865 | 0.010 | nCSF |
| *S_1246 | ADNI1 | 54.61 | 723.1 | 0.076 | aCSF |
| *S_1247 | ADNI1 | 31.94 | 649.1 | 0.049 | aCSF |
| *S_1250 | ADNI1 | 16.84 | 1643 | 0.010 | nCSF |
| *S_1260 | ADNI1 | 16.07 | 1470 | 0.011 | nCSF |
| *S_1262 | ADNI1 | 50.13 | 529.5 | 0.095 | aCSF |
| *S_1265 | ADNI1 | 26.61 | 860.9 | 0.031 | aCSF |
| *S_1268 | ADNI1 | 14.62 | 545.1 | 0.027 | nCSF |
| *S_1269 | ADNI1 | 46.2 | 997.6 | 0.046 | aCSF |
| *S_1281 | ADNI1 | 38.49 | 350.9 | 0.110 | aCSF |
| *S_1285 | ADNI1 | 74.51 | 862.9 | 0.086 | aCSF |
| *S_1293 | ADNI1 | 20.8 | 291.1 | 0.071 | aCSF |
| *S_1296 | ADNI1 | 24.37 | 318.8 | 0.076 | aCSF |
| *S_1309 | ADNI1 | 32.46 | 570 | 0.057 | aCSF |
| *S_1315 | ADNI1 | 34.36 | 761.5 | 0.045 | aCSF |
| *S_1318 | ADNI1 | 13.63 | 1134 | 0.012 | nCSF |
| *S_1321 | ADNI1 | 13.99 | 1113 | 0.013 | nCSF |
| *S_1341 | ADNI1 | 33.26 | 507.1 | 0.066 | aCSF |
| *S_1351 | ADNI1 | 58.26 | 755.5 | 0.077 | aCSF |
| *S_1352 | ADNI1 | 21.45 | 1685 | 0.013 | nCSF |
| *S_1371 | ADNI1 | 39.73 | 538.9 | 0.074 | aCSF |
| *S_1373 | ADNI1 | 29.77 | 430.7 | 0.069 | aCSF |
| *S_1379 | ADNI1 | 24.47 | 1539 | 0.016 | nCSF |
| *S_1394 | ADNI1 | 24.9 | 391.2 | 0.064 | aCSF |

| | | | | | |
|---|---|---|---|---|---|
| *S_1414 | ADNI1 | 18.62 | 1308 | 0.014 | nCSF |
| *S_1419 | ADNI1 | 22.48 | 390.5 | 0.058 | aCSF |
| *S_0166 | ADNI2 | 24.16 | 551.9 | 0.044 | aCSF |
| *S_0311 | ADNI2 | 21.32 | 1396 | 0.015 | nCSF |
| *S_0416 | ADNI2 | 17.14 | 1219 | 0.014 | nCSF |
| *S_0668 | ADNI2 | 15.31 | 1129 | 0.014 | nCSF |
| *S_1106 | ADNI2 | 27.39 | 829 | 0.033 | aCSF |
| *S_4001 | ADNI2 | 26.31 | 294.7 | 0.089 | aCSF |
| *S_4003 | ADNI2 | 28.47 | 1274 | 0.022 | nCSF |
| *S_4005 | ADNI2 | 36.63 | 853.5 | 0.043 | aCSF |
| *S_4007 | ADNI2 | 23.42 | 662.5 | 0.035 | aCSF |
| *S_4010 | ADNI2 | 14.52 | 681.8 | 0.021 | nCSF |
| *S_4012 | ADNI2 | 28.5 | 1009 | 0.028 | aCSF |
| *S_4014 | ADNI2 | 30.92 | 443.5 | 0.070 | aCSF |
| *S_4015 | ADNI2 | 39.81 | 384.6 | 0.104 | aCSF |
| *S_4018 | ADNI2 | 22.26 | 1893 | 0.012 | nCSF |
| *S_4020 | ADNI2 | 24.15 | 2559 | 0.009 | nCSF |
| *S_4021 | ADNI2 | 8 | 690.2 | 0.012 | nCSF |
| *S_4022 | ADNI2 | 47.72 | 427.9 | 0.112 | aCSF |
| *S_4024 | ADNI2 | 67.97 | 934.2 | 0.073 | aCSF |
| *S_4026 | ADNI2 | 18.33 | 509.2 | 0.036 | aCSF |
| *S_4028 | ADNI2 | 23.95 | 1910 | 0.013 | nCSF |
| *S_4029 | ADNI2 | 25.32 | 1063 | 0.024 | nCSF |
| *S_4030 | ADNI2 | 32.41 | 767.3 | 0.042 | aCSF |
| *S_4032 | ADNI2 | 12.23 | 1192 | 0.010 | nCSF |
| *S_4034 | ADNI2 | 46.46 | 486.2 | 0.096 | aCSF |
| *S_4035 | ADNI2 | 44.09 | 623.4 | 0.071 | aCSF |
| *S_4037 | ADNI2 | 17.76 | 1809 | 0.010 | nCSF |
| *S_4039 | ADNI2 | 57.04 | 593.6 | 0.096 | aCSF |
| *S_4042 | ADNI2 | 32.93 | 633.1 | 0.052 | aCSF |
| *S_4043 | ADNI2 | 21.98 | 2232 | 0.010 | nCSF |
| *S_4053 | ADNI2 | 50.45 | 879 | 0.057 | aCSF |
| *S_4057 | ADNI2 | 56.96 | 950.8 | 0.060 | aCSF |
| *S_4058 | ADNI2 | 28.03 | 434.5 | 0.065 | aCSF |
| *S_4059 | ADNI2 | 20.2 | 374.4 | 0.054 | aCSF |
| *S_4060 | ADNI2 | 14.75 | 737.5 | 0.020 | nCSF |
| *S_4067 | ADNI2 | 15.53 | 396.6 | 0.039 | aCSF |
| *S_4071 | ADNI2 | 33.3 | 701.3 | 0.047 | aCSF |
| *S_4073 | ADNI2 | 15.34 | 836.7 | 0.018 | nCSF |
| *S_4075 | ADNI2 | 25.94 | 2093 | 0.012 | nCSF |
| *S_4076 | ADNI2 | 15.14 | 1939 | 0.008 | nCSF |

| | | | | | |
|---|---|---|---|---|---|
| *S_4079 | ADNI2 | 50.27 | 704.5 | 0.071 | aCSF |
| *S_4080 | ADNI2 | 28.08 | 950.6 | 0.030 | aCSF |
| *S_4081 | ADNI2 | 22.37 | 715.5 | 0.031 | aCSF |
| *S_4082 | ADNI2 | 22.53 | 1616 | 0.014 | nCSF |
| *S_4084 | ADNI2 | 20.67 | 2349 | 0.009 | nCSF |
| *S_4089 | ADNI2 | 57.63 | 751.8 | 0.077 | aCSF |
| *S_4090 | ADNI2 | 17.53 | 1687 | 0.010 | nCSF |
| *S_4093 | ADNI2 | 10.42 | 858.7 | 0.012 | nCSF |
| *S_4096 | ADNI2 | 20.04 | 267.2 | 0.075 | aCSF |
| *S_4097 | ADNI2 | 21.89 | 801.7 | 0.027 | nCSF |
| *S_4100 | ADNI2 | 28.58 | 889.9 | 0.032 | aCSF |
| *S_4103 | ADNI2 | 17.69 | 1432 | 0.012 | nCSF |
| *S_4104 | ADNI2 | 13.66 | 917.4 | 0.015 | nCSF |
| *S_4105 | ADNI2 | 11.29 | 974.3 | 0.012 | nCSF |
| *S_4114 | ADNI2 | 47.82 | 916.1 | 0.052 | aCSF |
| *S_4119 | ADNI2 | 30.31 | 1826 | 0.017 | nCSF |
| *S_4120 | ADNI2 | 30.52 | 666.1 | 0.046 | aCSF |
| *S_4121 | ADNI2 | 26.15 | 1345 | 0.019 | nCSF |
| *S_4122 | ADNI2 | 36.58 | 1002 | 0.037 | aCSF |
| *S_4125 | ADNI2 | 14.88 | 1685 | 0.009 | nCSF |
| *S_4127 | ADNI2 | 8.21 | 744.5 | 0.011 | nCSF |
| *S_4128 | ADNI2 | 22.98 | 879.1 | 0.026 | nCSF |
| *S_4131 | ADNI2 | 28.22 | 589.1 | 0.048 | aCSF |
| *S_4134 | ADNI2 | 31.45 | 836.8 | 0.038 | aCSF |
| *S_4139 | ADNI2 | 10.84 | 1415 | 0.008 | nCSF |
| *S_4146 | ADNI2 | 19.83 | 519.9 | 0.038 | aCSF |
| *S_4148 | ADNI2 | 15.01 | 1352 | 0.011 | nCSF |
| *S_4149 | ADNI2 | 18.54 | 958.3 | 0.019 | nCSF |
| *S_4150 | ADNI2 | 19.39 | 2118 | 0.009 | nCSF |
| *S_4151 | ADNI2 | 17.85 | 492.7 | 0.036 | aCSF |
| *S_4153 | ADNI2 | 42.01 | 519.2 | 0.081 | aCSF |
| *S_4157 | ADNI2 | 29.16 | 1079 | 0.027 | nCSF |
| *S_4158 | ADNI2 | 16.87 | 1644 | 0.010 | nCSF |
| *S_4160 | ADNI2 | 10.84 | 1035 | 0.010 | nCSF |
| *S_4162 | ADNI2 | 35.37 | 778.8 | 0.045 | aCSF |
| *S_4164 | ADNI2 | 19.89 | 2326 | 0.009 | nCSF |
| *S_4167 | ADNI2 | 29.43 | 645.3 | 0.046 | aCSF |
| *S_4168 | ADNI2 | 21.1 | 723.4 | 0.029 | aCSF |
| *S_4170 | ADNI2 | 17.37 | 1073 | 0.016 | nCSF |
| *S_4171 | ADNI2 | 55 | 765.6 | 0.072 | aCSF |
| *S_4172 | ADNI2 | 47.67 | 651.4 | 0.073 | aCSF |

| | | | | | |
|---|---|---|---|---|---|
| *S_4173 | ADNI2 | 13.03 | 1358 | 0.010 | nCSF |
| *S_4174 | ADNI2 | 26.77 | 734.6 | 0.036 | aCSF |
| *S_4176 | ADNI2 | 46.98 | 1156 | 0.041 | aCSF |
| *S_4177 | ADNI2 | 15.28 | 924.7 | 0.017 | nCSF |
| *S_4179 | ADNI2 | 20.71 | 301.5 | 0.069 | aCSF |
| *S_4185 | ADNI2 | 9.55 | 723.1 | 0.013 | nCSF |
| *S_4187 | ADNI2 | 18.85 | 963.7 | 0.020 | nCSF |
| *S_4188 | ADNI2 | 27.59 | 485.5 | 0.057 | aCSF |
| *S_4189 | ADNI2 | 33.9 | 545.2 | 0.062 | aCSF |
| *S_4192 | ADNI2 | 21.54 | 564.5 | 0.038 | aCSF |
| *S_4195 | ADNI2 | 55.81 | 433.9 | 0.129 | aCSF |
| *S_4196 | ADNI2 | 36.31 | 1400 | 0.026 | nCSF |
| *S_4197 | ADNI2 | 17.74 | 555.4 | 0.032 | aCSF |
| *S_4198 | ADNI2 | 19.27 | 712.2 | 0.027 | nCSF |
| *S_4200 | ADNI2 | 13.05 | 1544 | 0.008 | nCSF |
| *S_4201 | ADNI2 | 59.94 | 749.4 | 0.080 | aCSF |
| *S_4205 | ADNI2 | 27.46 | 992.5 | 0.028 | nCSF |
| *S_4208 | ADNI2 | 16.95 | 1823 | 0.009 | nCSF |
| *S_4211 | ADNI2 | 20.36 | 456 | 0.045 | aCSF |
| *S_4213 | ADNI2 | 18.85 | 1691 | 0.011 | nCSF |
| *S_4214 | ADNI2 | 9.39 | 879.9 | 0.011 | nCSF |
| *S_4215 | ADNI2 | 27.32 | 576.4 | 0.047 | aCSF |
| *S_4219 | ADNI2 | 10.16 | 908.7 | 0.011 | nCSF |
| *S_4223 | ADNI2 | 23.49 | 834.6 | 0.028 | aCSF |
| *S_4224 | ADNI2 | 15.43 | 1201 | 0.013 | nCSF |
| *S_4225 | ADNI2 | 38.07 | 713.7 | 0.053 | aCSF |
| *S_4229 | ADNI2 | 9.92 | 925 | 0.011 | nCSF |
| *S_4232 | ADNI2 | 17.1 | 682 | 0.025 | nCSF |
| *S_4235 | ADNI2 | 32.64 | 697.2 | 0.047 | aCSF |
| *S_4240 | ADNI2 | 27.73 | 337.5 | 0.082 | aCSF |
| *S_4241 | ADNI2 | 32.12 | 731.9 | 0.044 | aCSF |
| *S_4243 | ADNI2 | 36.35 | 1069 | 0.034 | aCSF |
| *S_4244 | ADNI2 | 12.16 | 654.3 | 0.019 | nCSF |
| *S_4245 | ADNI2 | 9.22 | 924.4 | 0.010 | nCSF |
| *S_4250 | ADNI2 | 43.93 | 757.1 | 0.058 | aCSF |
| *S_4251 | ADNI2 | 21.73 | 728.5 | 0.030 | aCSF |
| *S_4252 | ADNI2 | 32.08 | 556.4 | 0.058 | aCSF |
| *S_4254 | ADNI2 | 31.96 | 1194 | 0.027 | nCSF |
| *S_4258 | ADNI2 | 34.52 | 641.6 | 0.054 | aCSF |
| *S_4259 | ADNI2 | 17.37 | 469.5 | 0.037 | aCSF |
| *S_4262 | ADNI2 | 34.28 | 790.5 | 0.043 | aCSF |

| | | | | | |
|---|---|---|---|---|---|
| *S_4263 | ADNI2 | 41.11 | 669.3 | 0.061 | aCSF |
| *S_4266 | ADNI2 | 33.22 | 820.1 | 0.041 | aCSF |
| *S_4269 | ADNI2 | 17.48 | 1157 | 0.015 | nCSF |
| *S_4270 | ADNI2 | 19.03 | 479.7 | 0.040 | aCSF |
| *S_4272 | ADNI2 | 20.08 | 601.8 | 0.033 | aCSF |
| *S_4274 | ADNI2 | 13.63 | 893.6 | 0.015 | nCSF |
| *S_4275 | ADNI2 | 20.11 | 1775 | 0.011 | nCSF |
| *S_4276 | ADNI2 | 21.92 | 2335 | 0.009 | nCSF |
| *S_4278 | ADNI2 | 19.33 | 422.8 | 0.046 | aCSF |
| *S_4279 | ADNI2 | 29.37 | 2612 | 0.011 | nCSF |
| *S_4287 | ADNI2 | 46.13 | 737.1 | 0.063 | aCSF |
| *S_4288 | ADNI2 | 10.96 | 716.6 | 0.015 | nCSF |
| *S_4290 | ADNI2 | 30.12 | 510.8 | 0.059 | aCSF |
| *S_4291 | ADNI2 | 18.84 | 762 | 0.025 | nCSF |
| *S_4292 | ADNI2 | 19.56 | 1635 | 0.012 | nCSF |
| *S_4294 | ADNI2 | 30.02 | 535 | 0.056 | aCSF |
| *S_4300 | ADNI2 | 14.1 | 633 | 0.022 | nCSF |
| *S_4301 | ADNI2 | 10.88 | 1052 | 0.010 | nCSF |
| *S_4302 | ADNI2 | 31.98 | 948.3 | 0.034 | aCSF |
| *S_4303 | ADNI2 | 24.99 | 668.7 | 0.037 | aCSF |
| *S_4308 | ADNI2 | 20.34 | 2040 | 0.010 | nCSF |
| *S_4310 | ADNI2 | 11.45 | 659.3 | 0.017 | nCSF |
| *S_4312 | ADNI2 | 20.75 | 666.2 | 0.031 | aCSF |
| *S_4313 | ADNI2 | 14.48 | 1244 | 0.012 | nCSF |
| *S_4320 | ADNI2 | 22 | 979.7 | 0.022 | nCSF |
| *S_4324 | ADNI2 | 15.05 | 342.6 | 0.044 | aCSF |
| *S_4331 | ADNI2 | 12.78 | 992 | 0.013 | nCSF |
| *S_4335 | ADNI2 | 16.11 | 203 | 0.079 | aCSF |
| *S_4337 | ADNI2 | 9.9 | 998.6 | 0.010 | nCSF |
| *S_4339 | ADNI2 | 27.13 | 481.2 | 0.056 | aCSF |
| *S_4340 | ADNI2 | 15.4 | 1913 | 0.008 | nCSF |
| *S_4343 | ADNI2 | 53.38 | 695.5 | 0.077 | aCSF |
| *S_4346 | ADNI2 | 26.35 | 664.5 | 0.040 | aCSF |
| *S_4348 | ADNI2 | 18.37 | 798.6 | 0.023 | nCSF |
| *S_4349 | ADNI2 | 17.82 | 995.8 | 0.018 | nCSF |
| *S_4350 | ADNI2 | 18.8 | 1918 | 0.010 | nCSF |
| *S_4351 | ADNI2 | 24.86 | 703.7 | 0.035 | aCSF |
| *S_4352 | ADNI2 | 23.04 | 1882 | 0.012 | nCSF |
| *S_4357 | ADNI2 | 19.25 | 1332 | 0.014 | nCSF |
| *S_4359 | ADNI2 | 46.74 | 871.9 | 0.054 | aCSF |
| *S_4363 | ADNI2 | 36.15 | 682.1 | 0.053 | aCSF |

| | | | | | |
|---|---|---|---|---|---|
| *S_4365 | ADNI2 | 17.29 | 1264 | 0.014 | nCSF |
| *S_4366 | ADNI2 | 46.64 | 729.9 | 0.064 | aCSF |
| *S_4367 | ADNI2 | 20.11 | 938.8 | 0.021 | nCSF |
| *S_4369 | ADNI2 | 24.56 | 2249 | 0.011 | nCSF |
| *S_4371 | ADNI2 | 24.15 | 957.2 | 0.025 | nCSF |
| *S_4376 | ADNI2 | 23.97 | 2108 | 0.011 | nCSF |
| *S_4382 | ADNI2 | 32.3 | 3088 | 0.010 | nCSF |
| *S_4384 | ADNI2 | 9.86 | 867.5 | 0.011 | nCSF |
| *S_4385 | ADNI2 | 12.76 | 220.6 | 0.058 | aCSF |
| *S_4386 | ADNI2 | 48.1 | 1221 | 0.039 | aCSF |
| *S_4387 | ADNI2 | 21.77 | 1099 | 0.020 | nCSF |
| *S_4388 | ADNI2 | 16.55 | 874.3 | 0.019 | nCSF |
| *S_4391 | ADNI2 | 13.94 | 1731 | 0.008 | nCSF |
| *S_4392 | ADNI2 | 37.26 | 976.6 | 0.038 | aCSF |
| *S_4393 | ADNI2 | 30.8 | 2573 | 0.012 | nCSF |
| *S_4394 | ADNI2 | 14.77 | 1086 | 0.014 | nCSF |
| *S_4395 | ADNI2 | 11.81 | 982.1 | 0.012 | nCSF |
| *S_4396 | ADNI2 | 13.69 | 981.6 | 0.014 | nCSF |
| *S_4399 | ADNI2 | 19.62 | 1554 | 0.013 | nCSF |
| *S_4401 | ADNI2 | 32.26 | 1867 | 0.017 | nCSF |
| *S_4402 | ADNI2 | 46.42 | 456.8 | 0.102 | aCSF |
| *S_4404 | ADNI2 | 21.19 | 652 | 0.033 | aCSF |
| *S_4405 | ADNI2 | 44.5 | 633.8 | 0.070 | aCSF |
| *S_4406 | ADNI2 | 25.55 | 810.6 | 0.032 | aCSF |
| *S_4408 | ADNI2 | 28.72 | 611.9 | 0.047 | aCSF |
| *S_4410 | ADNI2 | 29.14 | 2019 | 0.014 | nCSF |
| *S_4414 | ADNI2 | 43.22 | 969.7 | 0.045 | aCSF |
| *S_4415 | ADNI2 | 92.08 | 389 | 0.237 | aCSF |
| *S_4417 | ADNI2 | 19.6 | 801.2 | 0.024 | nCSF |
| *S_4420 | ADNI2 | 32.22 | 795 | 0.041 | aCSF |
| *S_4421 | ADNI2 | 17.55 | 1955 | 0.009 | nCSF |
| *S_4422 | ADNI2 | 30.51 | 1373 | 0.022 | nCSF |
| *S_4423 | ADNI2 | 19.7 | 741 | 0.027 | nCSF |
| *S_4424 | ADNI2 | 14.6 | 752.9 | 0.019 | nCSF |
| *S_4426 | ADNI2 | 17.89 | 819.6 | 0.022 | nCSF |
| *S_4427 | ADNI2 | 20.73 | 1689 | 0.012 | nCSF |
| *S_4428 | ADNI2 | 16.67 | 1454 | 0.011 | nCSF |
| *S_4429 | ADNI2 | 25.29 | 2175 | 0.012 | nCSF |
| *S_4430 | ADNI2 | 54.28 | 590.5 | 0.092 | aCSF |
| *S_4433 | ADNI2 | 33.76 | 486.5 | 0.069 | aCSF |
| *S_4438 | ADNI2 | 17.4 | 627.4 | 0.028 | nCSF |

| | | | | | |
|---|---|---|---|---|---|
| *S_4442 | ADNI2 | 51.24 | 425.6 | 0.120 | aCSF |
| *S_4444 | ADNI2 | 25.37 | 971.6 | 0.026 | nCSF |
| *S_4447 | ADNI2 | 45.51 | 871 | 0.052 | aCSF |
| *S_4448 | ADNI2 | 26.82 | 2426 | 0.011 | nCSF |
| *S_4449 | ADNI2 | 17.99 | 957.1 | 0.019 | nCSF |
| *S_4453 | ADNI2 | 19.29 | 2069 | 0.009 | nCSF |
| *S_4456 | ADNI2 | 27.26 | 689.2 | 0.040 | aCSF |
| *S_4458 | ADNI2 | 51.53 | 651.3 | 0.079 | aCSF |
| *S_4462 | ADNI2 | 21.12 | 721.6 | 0.029 | aCSF |
| *S_4464 | ADNI2 | 17.47 | 840.8 | 0.021 | nCSF |
| *S_4466 | ADNI2 | 15.52 | 1061 | 0.015 | nCSF |
| *S_4467 | ADNI2 | 76.51 | 705.2 | 0.108 | aCSF |
| *S_4469 | ADNI2 | 34.16 | 3384 | 0.010 | nCSF |
| *S_4473 | ADNI2 | 40.95 | 998.8 | 0.041 | aCSF |
| *S_4474 | ADNI2 | 36.17 | 507.2 | 0.071 | aCSF |
| *S_4477 | ADNI2 | 33.6 | 651.4 | 0.052 | aCSF |
| *S_4480 | ADNI2 | 23.17 | 682.1 | 0.034 | aCSF |
| *S_4482 | ADNI2 | 59.99 | 1244 | 0.048 | aCSF |
| *S_4483 | ADNI2 | 15.26 | 1696 | 0.009 | nCSF |
| *S_4485 | ADNI2 | 25.67 | 1563 | 0.016 | nCSF |
| *S_4488 | ADNI2 | 16.28 | 1641 | 0.010 | nCSF |
| *S_4491 | ADNI2 | 14.19 | 1292 | 0.011 | nCSF |
| *S_4494 | ADNI2 | 46.73 | 308.5 | 0.151 | aCSF |
| *S_4496 | ADNI2 | 28.25 | 3462 | 0.008 | nCSF |
| *S_4498 | ADNI2 | 35.75 | 876.6 | 0.041 | aCSF |
| *S_4499 | ADNI2 | 26.29 | 1841 | 0.014 | nCSF |
| *S_4500 | ADNI2 | 37.41 | 510.7 | 0.073 | aCSF |
| *S_4501 | ADNI2 | 31.57 | 741.5 | 0.043 | aCSF |
| *S_4502 | ADNI2 | 60.23 | 799.3 | 0.075 | aCSF |
| *S_4503 | ADNI2 | 17.1 | 1245 | 0.014 | nCSF |
| *S_4505 | ADNI2 | 28.49 | 2385 | 0.012 | nCSF |
| *S_4507 | ADNI2 | 30.48 | 697.3 | 0.044 | aCSF |
| *S_4508 | ADNI2 | 17.34 | 685.5 | 0.025 | nCSF |
| *S_4510 | ADNI2 | 47.52 | 733 | 0.065 | aCSF |
| *S_4513 | ADNI2 | 15.28 | 1085 | 0.014 | nCSF |
| *S_4515 | ADNI2 | 32.49 | 520.3 | 0.062 | aCSF |
| *S_4516 | ADNI2 | 19.91 | 1488 | 0.013 | nCSF |
| *S_4520 | ADNI2 | 35.59 | 2608 | 0.014 | nCSF |
| *S_4521 | ADNI2 | 50.4 | 918.5 | 0.055 | aCSF |
| *S_4522 | ADNI2 | 10.76 | 1084 | 0.010 | nCSF |
| *S_4526 | ADNI2 | 49.71 | 366 | 0.136 | aCSF |

| | | | | | |
|---|---|---|---|---|---|
| *S_4530 | ADNI2 | 19.88 | 592.3 | 0.034 | aCSF |
| *S_4531 | ADNI2 | 19.32 | 464.8 | 0.042 | aCSF |
| *S_4538 | ADNI2 | 28.11 | 761 | 0.037 | aCSF |
| *S_4539 | ADNI2 | 11.13 | 1093 | 0.010 | nCSF |
| *S_4542 | ADNI2 | 36.12 | 479.4 | 0.075 | aCSF |
| *S_4545 | ADNI2 | 11.89 | 1373 | 0.009 | nCSF |
| *S_4546 | ADNI2 | 24.81 | 523.9 | 0.047 | aCSF |
| *S_4547 | ADNI2 | 23.21 | 695.1 | 0.033 | aCSF |
| *S_4549 | ADNI2 | 32.02 | 531.8 | 0.060 | aCSF |
| *S_4552 | ADNI2 | 19.97 | 1241 | 0.016 | nCSF |
| *S_4559 | ADNI2 | 17.06 | 1969 | 0.009 | nCSF |
| *S_4560 | ADNI2 | 18.98 | 2140 | 0.009 | nCSF |
| *S_4562 | ADNI2 | 23.97 | 312.4 | 0.077 | aCSF |
| *S_4565 | ADNI2 | 16.72 | 495.8 | 0.034 | aCSF |
| *S_4568 | ADNI2 | 54.7 | 590.9 | 0.093 | aCSF |
| *S_4576 | ADNI2 | 55.14 | 2254 | 0.024 | nCSF |
| *S_4577 | ADNI2 | 12.84 | 318.3 | 0.040 | aCSF |
| *S_4578 | ADNI2 | 14.28 | 2054 | 0.007 | nCSF |
| *S_4580 | ADNI2 | 25.42 | 857 | 0.030 | aCSF |
| *S_4582 | ADNI2 | 33.56 | 786.9 | 0.043 | aCSF |
| *S_4584 | ADNI2 | 32.75 | 806 | 0.041 | aCSF |
| *S_4585 | ADNI2 | 18.76 | 1435 | 0.013 | nCSF |
| *S_4586 | ADNI2 | 19.26 | 1744 | 0.011 | nCSF |
| *S_4587 | ADNI2 | 24.38 | 686.4 | 0.036 | aCSF |
| *S_4589 | ADNI2 | 24.39 | 805.3 | 0.030 | aCSF |
| *S_4591 | ADNI2 | 28.67 | 651 | 0.044 | aCSF |
| *S_4595 | ADNI2 | 53 | 957.2 | 0.055 | aCSF |
| *S_4596 | ADNI2 | 23.72 | 473.8 | 0.050 | aCSF |
| *S_4597 | ADNI2 | 27.88 | 715.5 | 0.039 | aCSF |
| *S_4598 | ADNI2 | 29.77 | 2696 | 0.011 | nCSF |
| *S_4599 | ADNI2 | 15.8 | 772.8 | 0.020 | nCSF |
| *S_4604 | ADNI2 | 14.03 | 1279 | 0.011 | nCSF |
| *S_4605 | ADNI2 | 28.01 | 817.7 | 0.034 | aCSF |
| *S_4611 | ADNI2 | 20.44 | 574.2 | 0.036 | aCSF |
| *S_4612 | ADNI2 | 13.89 | 499.5 | 0.028 | nCSF |
| *S_4615 | ADNI2 | 30.62 | 864.5 | 0.035 | aCSF |
| *S_4616 | ADNI2 | 20.49 | 897.8 | 0.023 | nCSF |
| *S_4621 | ADNI2 | 14.55 | 853.3 | 0.017 | nCSF |
| *S_4623 | ADNI2 | 20.08 | 635.2 | 0.032 | aCSF |
| *S_4624 | ADNI2 | 37.45 | 608.9 | 0.062 | aCSF |
| *S_4625 | ADNI2 | 36.69 | 366.3 | 0.100 | aCSF |

| | | | | | |
|---|---|---|---|---|---|
| *S_4629 | ADNI2 | 10.33 | 1088 | 0.009 | nCSF |
| *S_4631 | ADNI2 | 30.41 | 725.4 | 0.042 | aCSF |
| *S_4635 | ADNI2 | 20.45 | 359.3 | 0.057 | aCSF |
| *S_4636 | ADNI2 | 30.4 | 744.4 | 0.041 | aCSF |
| *S_4637 | ADNI2 | 19.09 | 351.4 | 0.054 | aCSF |
| *S_4638 | ADNI2 | 15.52 | 1739 | 0.009 | nCSF |
| *S_4641 | ADNI2 | 21.43 | 660.7 | 0.032 | aCSF |
| *S_4643 | ADNI2 | 16.16 | 1704 | 0.009 | nCSF |
| *S_4644 | ADNI2 | 17.36 | 1778 | 0.010 | nCSF |
| *S_4645 | ADNI2 | 21.56 | 539.1 | 0.040 | aCSF |
| *S_4657 | ADNI2 | 59.14 | 883.8 | 0.067 | aCSF |
| *S_4659 | ADNI2 | 27.55 | 667.9 | 0.041 | aCSF |
| *S_4660 | ADNI2 | 62.39 | 767.6 | 0.081 | aCSF |
| *S_4661 | ADNI2 | 33.6 | 604 | 0.056 | aCSF |
| *S_4668 | ADNI2 | 33.14 | 698.1 | 0.047 | aCSF |
| *S_4672 | ADNI2 | 33.17 | 447.8 | 0.074 | aCSF |
| *S_4674 | ADNI2 | 31.89 | 843.1 | 0.038 | aCSF |
| *S_4675 | ADNI2 | 52.77 | 733.7 | 0.072 | aCSF |
| *S_4679 | ADNI2 | 31.59 | 1046 | 0.030 | aCSF |
| *S_4680 | ADNI2 | 24.13 | 487.9 | 0.049 | aCSF |
| *S_4686 | ADNI2 | 36.78 | 769.7 | 0.048 | aCSF |
| *S_4688 | ADNI2 | 31.12 | 1003 | 0.031 | aCSF |
| *S_4689 | ADNI2 | 32.4 | 747.3 | 0.043 | aCSF |
| *S_4696 | ADNI2 | 46.11 | 541.8 | 0.085 | aCSF |
| *S_4707 | ADNI2 | 17.93 | 451.7 | 0.040 | aCSF |
| *S_4713 | ADNI2 | 26.7 | 954.8 | 0.028 | nCSF |
| *S_4714 | ADNI2 | 36.67 | 915.7 | 0.040 | aCSF |
| *S_4715 | ADNI2 | 64.5 | 1009 | 0.064 | aCSF |
| *S_4718 | ADNI2 | 29.81 | 411.7 | 0.072 | aCSF |
| *S_4720 | ADNI2 | 31.28 | 483.9 | 0.065 | aCSF |
| *S_4721 | ADNI2 | 15.52 | 746.4 | 0.021 | nCSF |
| *S_4728 | ADNI2 | 31.27 | 507 | 0.062 | aCSF |
| *S_4729 | ADNI2 | 74.27 | 748.8 | 0.099 | aCSF |
| *S_4730 | ADNI2 | 24.06 | 588.2 | 0.041 | aCSF |
| *S_4732 | ADNI2 | 38.42 | 402.4 | 0.095 | aCSF |
| *S_4736 | ADNI2 | 28.48 | 440.3 | 0.065 | aCSF |
| *S_4739 | ADNI2 | 14.68 | 1404 | 0.010 | nCSF |
| *S_4742 | ADNI2 | 8.94 | 852.7 | 0.010 | nCSF |
| *S_4743 | ADNI2 | 22.83 | 844.8 | 0.027 | nCSF |
| *S_4744 | ADNI2 | 57.71 | 617.8 | 0.093 | aCSF |
| *S_4746 | ADNI2 | 20.32 | 880.2 | 0.023 | nCSF |

| | | | | | |
|---|---|---|---|---|---|
| *S_4756 | ADNI2 | 27.12 | 675.3 | 0.040 | aCSF |
| *S_4757 | ADNI2 | 59.81 | 701.5 | 0.085 | aCSF |
| *S_4762 | ADNI2 | 9.56 | 1020 | 0.009 | nCSF |
| *S_4764 | ADNI2 | 27.57 | 912.4 | 0.030 | aCSF |
| *S_4765 | ADNI2 | 50.75 | 845.8 | 0.060 | aCSF |
| *S_4767 | ADNI2 | 17.02 | 822.1 | 0.021 | nCSF |
| *S_4770 | ADNI2 | 24.93 | 503.1 | 0.050 | aCSF |
| *S_4774 | ADNI2 | 71.75 | 877.3 | 0.082 | aCSF |
| *S_4782 | ADNI2 | 20.02 | 312.3 | 0.064 | aCSF |
| *S_4783 | ADNI2 | 22.07 | 596.3 | 0.037 | aCSF |
| *S_4793 | ADNI2 | 30.19 | 443.9 | 0.068 | aCSF |
| *S_4795 | ADNI2 | 9.91 | 978.2 | 0.010 | nCSF |
| *S_4796 | ADNI2 | 74.45 | 755.2 | 0.099 | aCSF |
| *S_4799 | ADNI2 | 21.53 | 779.1 | 0.028 | nCSF |
| *S_4801 | ADNI2 | 26.96 | 375.7 | 0.072 | aCSF |
| *S_4802 | ADNI2 | 39.39 | 687.3 | 0.057 | aCSF |
| *S_4804 | ADNI2 | 14.99 | 432.8 | 0.035 | aCSF |
| *S_4805 | ADNI2 | 46 | 925 | 0.050 | aCSF |
| *S_4806 | ADNI2 | 8.87 | 927.5 | 0.010 | nCSF |
| *S_4807 | ADNI2 | 86.51 | 991.8 | 0.087 | aCSF |
| *S_4809 | ADNI2 | 26.79 | 782.6 | 0.034 | aCSF |
| *S_4815 | ADNI2 | 60.12 | 781.3 | 0.077 | aCSF |
| *S_4816 | ADNI2 | 52.08 | 511.3 | 0.102 | aCSF |
| *S_4817 | ADNI2 | 11.94 | 600.2 | 0.020 | nCSF |
| *S_4820 | ADNI2 | 64.84 | 384.4 | 0.169 | aCSF |
| *S_4832 | ADNI2 | 12.92 | 1236 | 0.010 | nCSF |
| *S_4835 | ADNI2 | 26.62 | 728.6 | 0.037 | aCSF |
| *S_4845 | ADNI2 | 38.18 | 652.1 | 0.059 | aCSF |
| *S_4849 | ADNI2 | 26.9 | 691.3 | 0.039 | aCSF |
| *S_4852 | ADNI2 | 58.19 | 685.7 | 0.085 | aCSF |
| *S_4853 | ADNI2 | 83.3 | 691.9 | 0.120 | aCSF |
| *S_4855 | ADNI2 | 21.48 | 732.8 | 0.029 | aCSF |
| *S_4856 | ADNI2 | 10.01 | 975.8 | 0.010 | nCSF |
| *S_4857 | ADNI2 | 36.33 | 888.5 | 0.041 | aCSF |
| *S_4858 | ADNI2 | 42.25 | 662.4 | 0.064 | aCSF |
| *S_4863 | ADNI2 | 32.73 | 449.4 | 0.073 | aCSF |
| *S_4867 | ADNI2 | 37.72 | 624.1 | 0.060 | aCSF |
| *S_4868 | ADNI2 | 30.84 | 519.7 | 0.059 | aCSF |
| *S_4877 | ADNI2 | 34.33 | 534.4 | 0.064 | aCSF |
| *S_4878 | ADNI2 | 14.75 | 1768 | 0.008 | nCSF |
| *S_4879 | ADNI2 | 37.91 | 532.9 | 0.071 | aCSF |

| | | | | | |
|---|---|---|---|---|---|
| *S_4885 | ADNI2 | 56.81 | 408.4 | 0.139 | aCSF |
| *S_4888 | ADNI2 | 31 | 876.8 | 0.035 | aCSF |
| *S_4889 | ADNI2 | 9.84 | 956.2 | 0.010 | nCSF |
| *S_4891 | ADNI2 | 25.35 | 533.5 | 0.048 | aCSF |
| *S_4893 | ADNI2 | 39.87 | 784.3 | 0.051 | aCSF |
| *S_4894 | ADNI2 | 47.59 | 423.6 | 0.112 | aCSF |
| *S_4897 | ADNI2 | 18.42 | 837.7 | 0.022 | nCSF |
| *S_4898 | ADNI2 | 16.42 | 711.2 | 0.023 | nCSF |
| *S_4900 | ADNI2 | 14.92 | 1605 | 0.009 | nCSF |
| *S_4902 | ADNI2 | 29.74 | 818.6 | 0.036 | aCSF |
| *S_4903 | ADNI2 | 27.5 | 854.8 | 0.032 | aCSF |
| *S_4904 | ADNI2 | 24.16 | 646.7 | 0.037 | aCSF |
| *S_4905 | ADNI2 | 46.04 | 568.8 | 0.081 | aCSF |
| *S_4910 | ADNI2 | 29.25 | 749.9 | 0.039 | aCSF |
| *S_4911 | ADNI2 | 17.17 | 330.9 | 0.052 | aCSF |
| *S_4918 | ADNI2 | 44.3 | 584.2 | 0.076 | aCSF |
| *S_4919 | ADNI2 | 17.85 | 989.8 | 0.018 | nCSF |
| *S_4920 | ADNI2 | 19.81 | 597.7 | 0.033 | aCSF |
| *S_4921 | ADNI2 | 11.77 | 1291 | 0.009 | nCSF |
| *S_4925 | ADNI2 | 27.16 | 694 | 0.039 | aCSF |
| *S_4926 | ADNI2 | 10.46 | 799.5 | 0.013 | nCSF |
| *S_4928 | ADNI2 | 33.02 | 883.9 | 0.037 | aCSF |
| *S_4929 | ADNI2 | 45.24 | 563.2 | 0.080 | aCSF |
| *S_4936 | ADNI2 | 18.52 | 518.3 | 0.036 | aCSF |
| *S_4938 | ADNI2 | 25.04 | 664.4 | 0.038 | aCSF |
| *S_4940 | ADNI2 | 30.36 | 652 | 0.047 | aCSF |
| *S_4944 | ADNI2 | 19.22 | 664.8 | 0.029 | aCSF |
| *S_4945 | ADNI2 | 28.27 | 757 | 0.037 | aCSF |
| *S_4947 | ADNI2 | 31.48 | 574.7 | 0.055 | aCSF |
| *S_4949 | ADNI2 | 36.03 | 506.1 | 0.071 | aCSF |
| *S_4951 | ADNI2 | 12.12 | 1024 | 0.012 | nCSF |
| *S_4952 | ADNI2 | 17.18 | 2429 | 0.007 | nCSF |
| *S_4954 | ADNI2 | 24.52 | 816.4 | 0.030 | aCSF |
| *S_4955 | ADNI2 | 13.28 | 395 | 0.034 | aCSF |
| *S_4959 | ADNI2 | 19.02 | 349.8 | 0.054 | aCSF |
| *S_4962 | ADNI2 | 47.15 | 998.2 | 0.047 | aCSF |
| *S_4964 | ADNI2 | 39.35 | 549.7 | 0.072 | aCSF |
| *S_4971 | ADNI2 | 28.66 | 471.3 | 0.061 | aCSF |
| *S_4974 | ADNI2 | 69.87 | 677.4 | 0.103 | aCSF |
| *S_4976 | ADNI2 | 62.4 | 905.6 | 0.069 | aCSF |
| *S_4980 | ADNI2 | 16.06 | 449.8 | 0.036 | aCSF |

| | | | | | |
|---|---|---|---|---|---|
| *S_4982 | ADNI2 | 40.66 | 692 | 0.059 | aCSF |
| *S_4984 | ADNI2 | 38.54 | 750.3 | 0.051 | aCSF |
| *S_4986 | ADNI2 | 11.27 | 780.5 | 0.014 | nCSF |
| *S_4987 | ADNI2 | 19.83 | 1091 | 0.018 | nCSF |
| *S_4989 | ADNI2 | 9.02 | 1087 | 0.008 | nCSF |
| *S_4990 | ADNI2 | 42 | 593.9 | 0.071 | aCSF |
| *S_4992 | ADNI2 | 18.42 | 418 | 0.044 | aCSF |
| *S_5005 | ADNI2 | 25.69 | 318.1 | 0.081 | aCSF |
| *S_5006 | ADNI2 | 66.53 | 653.9 | 0.102 | aCSF |
| *S_5007 | ADNI2 | 14.7 | 622 | 0.024 | nCSF |
| *S_5012 | ADNI2 | 33.95 | 458.3 | 0.074 | aCSF |
| *S_5013 | ADNI2 | 22.72 | 454.5 | 0.050 | aCSF |
| *S_5014 | ADNI2 | 13.24 | 585.6 | 0.023 | nCSF |
| *S_5015 | ADNI2 | 28.64 | 783.5 | 0.037 | aCSF |
| *S_5017 | ADNI2 | 54.78 | 622.5 | 0.088 | aCSF |
| *S_5018 | ADNI2 | 76.16 | 489.4 | 0.156 | aCSF |
| *S_5019 | ADNI2 | 36.63 | 587.9 | 0.062 | aCSF |
| *S_5023 | ADNI2 | 23.3 | 2081 | 0.011 | nCSF |
| *S_5027 | ADNI2 | 32.67 | 493.4 | 0.066 | aCSF |
| *S_5028 | ADNI2 | 49.99 | 437.6 | 0.114 | aCSF |
| *S_5029 | ADNI2 | 68.07 | 277.5 | 0.245 | aCSF |
| *S_5037 | ADNI2 | 25.91 | 700.9 | 0.037 | aCSF |
| *S_5040 | ADNI2 | 17.12 | 1879 | 0.009 | nCSF |
| *S_5047 | ADNI2 | 36.08 | 1001 | 0.036 | aCSF |
| *S_5054 | ADNI2 | 38.22 | 785.8 | 0.049 | aCSF |
| *S_5058 | ADNI2 | 14.76 | 255.5 | 0.058 | aCSF |
| *S_5059 | ADNI2 | 30.02 | 453.7 | 0.066 | aCSF |
| *S_5062 | ADNI2 | 74.14 | 542.4 | 0.137 | aCSF |
| *S_5063 | ADNI2 | 40.94 | 551.8 | 0.074 | aCSF |
| *S_5070 | ADNI2 | 19.45 | 489.7 | 0.040 | aCSF |
| *S_5078 | ADNI2 | 11.44 | 1321 | 0.009 | nCSF |
| *S_5079 | ADNI2 | 21.78 | 2667 | 0.008 | nCSF |
| *S_5082 | ADNI2 | 13.68 | 1600 | 0.009 | nCSF |
| *S_5083 | ADNI2 | 15.92 | 1002 | 0.016 | nCSF |
| *S_5087 | ADNI2 | 41.32 | 639.1 | 0.065 | aCSF |
| *S_5091 | ADNI2 | 24.08 | 1551 | 0.016 | nCSF |
| *S_5093 | ADNI2 | 21.86 | 2060 | 0.011 | nCSF |
| *S_5095 | ADNI2 | 34.29 | 382.5 | 0.090 | aCSF |
| *S_5096 | ADNI2 | 23.73 | 613.7 | 0.039 | aCSF |
| *S_5100 | ADNI2 | 12.93 | 1536 | 0.008 | nCSF |
| *S_5102 | ADNI2 | 24.23 | 1406 | 0.017 | nCSF |

| | | | | | |
|---|---|---|---|---|---|
| *S_5109 | ADNI2 | 17.63 | 771.9 | 0.023 | nCSF |
| *S_5110 | ADNI2 | 14 | 2008 | 0.007 | nCSF |
| *S_5113 | ADNI2 | 17.65 | 1762 | 0.010 | nCSF |
| *S_5118 | ADNI2 | 12.16 | 1210 | 0.010 | nCSF |
| *S_5119 | ADNI2 | 74.52 | 910.3 | 0.082 | aCSF |
| *S_5124 | ADNI2 | 30.81 | 2705 | 0.011 | nCSF |
| *S_5125 | ADNI2 | 14.06 | 1518 | 0.009 | nCSF |
| *S_5127 | ADNI2 | 21.87 | 469.2 | 0.047 | aCSF |
| *S_5130 | ADNI2 | 13.47 | 1687 | 0.008 | nCSF |
| *S_5141 | ADNI2 | 18.57 | 1923 | 0.010 | nCSF |
| *S_5142 | ADNI2 | 30.29 | 849.9 | 0.036 | aCSF |
| *S_5147 | ADNI2 | 26.23 | 2124 | 0.012 | nCSF |
| *S_5153 | ADNI2 | 20.68 | 2239 | 0.009 | nCSF |
| *S_5157 | ADNI2 | 15.83 | 1666 | 0.010 | nCSF |
| *S_5169 | ADNI2 | 26.06 | 1490 | 0.017 | nCSF |
| *S_5178 | ADNI2 | 20.96 | 1623 | 0.013 | nCSF |
| *S_5185 | ADNI2 | 45.64 | 662.8 | 0.069 | aCSF |
| *S_5195 | ADNI2 | 32.99 | 2571 | 0.013 | nCSF |
| *S_5197 | ADNI2 | 13.05 | 1142 | 0.011 | nCSF |
| *S_5200 | ADNI2 | 9.63 | 1112 | 0.009 | nCSF |
| *S_5202 | ADNI2 | 21.4 | 545.3 | 0.039 | aCSF |
| *S_5213 | ADNI2 | 25.87 | 3012 | 0.009 | nCSF |
| *S_5219 | ADNI2 | 28.83 | 1145 | 0.025 | nCSF |
| *S_5228 | ADNI2 | 21.07 | 1709 | 0.012 | nCSF |
| *S_5230 | ADNI2 | 22.84 | 1006 | 0.023 | nCSF |
| *S_5242 | ADNI2 | 27.06 | 1306 | 0.021 | nCSF |
| *S_5243 | ADNI2 | 14.2 | 1528 | 0.009 | nCSF |
| *S_5252 | ADNI2 | 27.87 | 474.9 | 0.059 | aCSF |
| *S_5256 | ADNI2 | 36.38 | 923.3 | 0.039 | aCSF |
| *S_5258 | ADNI2 | 26.32 | 869.7 | 0.030 | aCSF |
| *S_5265 | ADNI2 | 24.78 | 466.1 | 0.053 | aCSF |
| *S_5266 | ADNI2 | 24.28 | 2433 | 0.010 | nCSF |
| *S_5272 | ADNI2 | 25.05 | 2358 | 0.011 | nCSF |
| *S_5277 | ADNI2 | 34.39 | 780.3 | 0.044 | aCSF |
| *S_5280 | ADNI2 | 10.08 | 731.8 | 0.014 | nCSF |
| *S_5287 | ADNI2 | 26.11 | 2358 | 0.011 | nCSF |
| *S_5289 | ADNI2 | 15.79 | 1665 | 0.009 | nCSF |
| *S_5292 | ADNI2 | 36.39 | 928.3 | 0.039 | aCSF |
| *S_5296 | ADNI2 | 20.9 | 1460 | 0.014 | nCSF |
| *S_0056 | ADNIGO | 15.18 | 1198 | 0.013 | nCSF |
| *S_0059 | ADNIGO | 16.22 | 1616 | 0.010 | nCSF |

| | | | | | |
|---|---|---|---|---|---|
| *S_0227 | ADNIGO | 28.45 | 323 | 0.088 | aCSF |
| *S_0272 | ADNIGO | 23.79 | 1431 | 0.017 | nCSF |
| *S_1030 | ADNIGO | 40.35 | 985.8 | 0.041 | aCSF |
| *S_1117 | ADNIGO | 52.7 | 329.2 | 0.160 | aCSF |
| *S_1261 | ADNIGO | 21.23 | 1711 | 0.012 | nCSF |
| *S_2022 | ADNIGO | 22.11 | 929.2 | 0.024 | nCSF |
| *S_2026 | ADNIGO | 9.42 | 1079 | 0.009 | nCSF |
| *S_2042 | ADNIGO | 10.58 | 827.6 | 0.013 | nCSF |
| *S_2045 | ADNIGO | 20.5 | 725.1 | 0.028 | aCSF |
| *S_2047 | ADNIGO | 40.95 | 709.2 | 0.058 | aCSF |
| *S_2055 | ADNIGO | 45.44 | 637.1 | 0.071 | aCSF |
| *S_2063 | ADNIGO | 29.1 | 564.4 | 0.052 | aCSF |
| *S_2068 | ADNIGO | 32.76 | 722.8 | 0.045 | aCSF |
| *S_2073 | ADNIGO | 24.09 | 1094 | 0.022 | nCSF |
| *S_2077 | ADNIGO | 18.71 | 859.1 | 0.022 | nCSF |
| *S_2079 | ADNIGO | 48.48 | 518.6 | 0.093 | aCSF |
| *S_2083 | ADNIGO | 13.69 | 1033 | 0.013 | nCSF |
| *S_2087 | ADNIGO | 37.46 | 436.7 | 0.086 | aCSF |
| *S_2100 | ADNIGO | 40.16 | 673.7 | 0.060 | aCSF |
| *S_2106 | ADNIGO | 48.31 | 361.3 | 0.134 | aCSF |
| *S_2109 | ADNIGO | 28.79 | 532.1 | 0.054 | aCSF |
| *S_2121 | ADNIGO | 20.87 | 1016 | 0.021 | nCSF |
| *S_2125 | ADNIGO | 15.85 | 638.6 | 0.025 | nCSF |
| *S_2130 | ADNIGO | 30.39 | 476.6 | 0.064 | aCSF |
| *S_2133 | ADNIGO | 28.71 | 413.9 | 0.069 | aCSF |
| *S_2142 | ADNIGO | 32.49 | 632.1 | 0.051 | aCSF |
| *S_2150 | ADNIGO | 24.9 | 731.4 | 0.034 | aCSF |
| *S_2153 | ADNIGO | 10.4 | 980.4 | 0.011 | nCSF |
| *S_2155 | ADNIGO | 82.04 | 539.6 | 0.152 | aCSF |
| *S_2164 | ADNIGO | 12.57 | 1097 | 0.011 | nCSF |
| *S_2167 | ADNIGO | 33.28 | 832 | 0.040 | aCSF |
| *S_2171 | ADNIGO | 42.04 | 738.2 | 0.057 | aCSF |
| *S_2190 | ADNIGO | 35.48 | 963.2 | 0.037 | aCSF |
| *S_2194 | ADNIGO | 42.38 | 884.2 | 0.048 | aCSF |
| *S_2195 | ADNIGO | 49.19 | 796 | 0.062 | aCSF |
| *S_2196 | ADNIGO | 22.11 | 695.3 | 0.032 | aCSF |
| *S_2201 | ADNIGO | 18.14 | 2282 | 0.008 | nCSF |
| *S_2205 | ADNIGO | 23.28 | 336.6 | 0.069 | aCSF |
| *S_2210 | ADNIGO | 11.79 | 316.1 | 0.037 | aCSF |
| *S_2216 | ADNIGO | 23.14 | 530 | 0.044 | aCSF |
| *S_2247 | ADNIGO | 14.24 | 858.9 | 0.017 | nCSF |

| | | | | | |
|---|---|---|---|---|---|
| *S_2248 | ADNIGO | 66.72 | 646.9 | 0.103 | aCSF |
| *S_2263 | ADNIGO | 13.19 | 776.9 | 0.017 | nCSF |
| *S_2264 | ADNIGO | 21.16 | 839.5 | 0.025 | nCSF |
| *S_2274 | ADNIGO | 25.43 | 476.5 | 0.053 | aCSF |
| *S_2284 | ADNIGO | 15.13 | 791.6 | 0.019 | nCSF |
| *S_2307 | ADNIGO | 32.58 | 741.3 | 0.044 | aCSF |
| *S_2308 | ADNIGO | 11.78 | 982.6 | 0.012 | nCSF |
| *S_2316 | ADNIGO | 69.57 | 589.6 | 0.118 | aCSF |
| *S_2324 | ADNIGO | 9.8 | 832.9 | 0.012 | nCSF |
| *S_2333 | ADNIGO | 23.45 | 662.6 | 0.035 | aCSF |
| *S_2336 | ADNIGO | 33.64 | 909.4 | 0.037 | aCSF |
| *S_2347 | ADNIGO | 24.18 | 1010 | 0.024 | nCSF |
| *S_2367 | ADNIGO | 18.85 | 1086 | 0.017 | nCSF |
| *S_2373 | ADNIGO | 68.61 | 779.8 | 0.088 | aCSF |
| *S_2379 | ADNIGO | 10.58 | 804.8 | 0.013 | nCSF |
| *S_2380 | ADNIGO | 21.81 | 696.5 | 0.031 | aCSF |
| *S_2381 | ADNIGO | 57.7 | 684.7 | 0.084 | aCSF |
| *S_2389 | ADNIGO | 11.73 | 881 | 0.013 | nCSF |
| *S_2390 | ADNIGO | 38.61 | 892.1 | 0.043 | aCSF |
| *S_2391 | ADNIGO | 19.78 | 412.7 | 0.048 | aCSF |
| *S_2394 | ADNIGO | 17.5 | 360.4 | 0.049 | aCSF |
| *S_2403 | ADNIGO | 24.66 | 572.4 | 0.043 | aCSF |

RID = ADNI Research Identification Number; pTau = cerebrospinal fluid (CSF) hyper phosphorylated tau; $A\beta_{1\text{-}42}$ = CSF measure of amyloid Beta (1-42 peptides), pTau/$A\beta_{1\text{-}42}$ = ratio of the hyperphsphorylated tau and amyloid Beta; Group CSF = CSF grouping strategy based on the pTau/ $A\beta_{1\text{-}42}$ ratio: aCSF = pTau/ $A\beta_{1\text{-}42}$ >= 0.028; nCSF = pTau/ $A\beta_{1\text{-}42}$ < 0.028. Aβ1-42 and pTau concentrations are in picograms per milliliter (pg/mL).

### References


Schmitz TW, Spreng RN. Basal forebrain degeneration precedes and predicts the cortical spread of Alzheimer's pathology. Nat Commun 2016; 7
